# Benchmarking long-read RNA sequencing for de novo transcriptome assembly in non-model plant species: insights from *Moricandia arvensis*

**DOI:** 10.64898/2026.09.08.750093

**Authors:** Saloni Sharma, Michael Hackenberg, Luis Navarro, Eduardo Narbona, Adela González-Megias, Cristina Armas, Jose M. Gomez, Francisco Perfectti

## Abstract

De novo transcriptome assembly is the standard approach for constructing a reference transcriptome in non-model plants that lack a high-quality genome, yet short-read assemblies struggle to resolve full-length isoforms. Long-read Iso-Seq (PacBio) captures full-length transcripts directly, but its use as a primary reference and the choice of downstream assembly pipeline remains poorly benchmarked. Here, we construct genome-free Iso-Seq reference transcriptomes for two organs, flower and leaf, of the non-model species *Moricandia arvensis* (L.) DC. (Brassicaceae), and systematically compare pipeline strategies combining Iso-Seq clustering, CD-HIT redundancy reduction, and Cogent graph-based reconstruction, benchmarked by BUSCO completeness, RSEM short-read mapping, and TransDecoder ORF completeness. We find that the optimal pipeline is organ specific. For flower, CD-HIT pre-filtering followed by Cogent reconstruction produced a high-quality reference (95.3% BUSCO complete). For the leaf, the same Cogent step was detrimental, reducing BUSCO completeness from 90.1% to 78.0% by incorrectly merging distinct genes; therefore, CD-HIT at 95% identity without reconstruction was retained. We trace this divergence to organ-specific input-data characteristics: leaf transcripts show extreme full-length-read expression skew and predominantly single-isoform gene support, depriving Cogent’s graph algorithm of the multi-isoform evidence it requires. We find that the concentration of full-length reads among the most highly expressed transcripts predicts pipeline suitability before reconstruction, with per-transcript read depth acting as a necessary but non-discriminating floor. Because the leaf reference lacked gene-level structure, we further recovered gene-isoform grouping using expression-aware read-clustering (Corset), which preserved completeness while restoring the paralog structure expected of a paleopolyploid genome and outperformed sequence-only clustering. Our results provide a robust, genome-free framework for constructing full-length reference transcriptomes in non-model plant species and demonstrate that pipeline choice must be evaluated per organ rather than assuming one size fits all.

## INTRODUCTION

Transcriptomics provides a formidable tool for elucidating the molecular mechanisms by which environmental stimuli modulate gene expression (Wang et al., 2009; Lockwood et al., 2015; Mante et al. 2025). The rapid advancement of high-throughput sequencing platforms and the broader integration of multi-omics approaches have substantially lowered the technical and economic barriers to transcriptomic analysis, enabling the precise quantification of transcript abundance and the characterization of alternative splicing events at a genome-wide scale (Manzoni et al., 2018). In non-model organisms, transcriptomics has proven particularly valuable for uncovering the molecular basis of ecologically relevant traits, including responses to biotic and abiotic stresses; however, the absence of a reference genome sequence introduces considerable analytical challenges that must be addressed through de novo assembly strategies (Clark, 2022; Kulaeva et al., 2017).

In the absence of a well-annotated, high-quality reference genome, de novo transcriptome assembly represents the standard approach for reconstructing the full complement of expressed transcripts from RNA-Seq data (Haas et al., 2013). While this strategy enables the identification of novel transcripts, it frequently yields fragmented assemblies that incompletely capture the true transcriptome (Martin and Wang, 2011). When available, genomic resources from phylogenetically related species may facilitate contig scaffolding and annotation; however, high-quality reference genomes remain scarce for many plant lineages, particularly those characterised by complex or highly diversified evolutionary histories (Soltis and Soltis, 2021; Marks et al., 2021). The generation of a comprehensive and accurate transcriptome resource is therefore of paramount importance in non-model plant species, as it underpins the reliable quantification of gene expression and the characterisation of regulatory networks. In this context, short-read RNA sequencing -despite its widespread adoption-is fundamentally limited in its capacity to resolve full-length isoforms, often producing chimeric or truncated transcript reconstructions that compromise the accuracy of downstream analyses and cross-species comparative transcriptomics (Byrne et al., 2019; Ament et al., 2025).

Single-molecule long-read Iso-Seq (PacBio) addresses many of these challenges by capturing full-length cDNAs and enabling direct transcript structure identification (Sharon et al., 2013; Shi et al., 2024), as it eliminates the requirement for transcript assembly and allows for fresh discoveries in a variety of applications (Weirather et al., 2017). An early landmark application of this approach in a non-model plant species was demonstrated by Li et al. (2017), who reconstructed full-length transcriptomes from two tissues of *Astragalus membranaceus* without a reference genome using PacBio Iso-Seq, providing a proof-of-concept pipeline for genome-free transcriptome construction in plants. Additionally, Iso-Seq is also increasingly used for annotation and isoform discovery; however, its application as a primary reference for de novo transcriptomics still requires further research (Wang et al., 2016; Cui et al., 2020; Shi et al., 2024). The present study builds upon and substantially extends Li et al. (2017) framework by systematically benchmarking multiple pipeline strategies across two organs, revealing critical organ-specific limitations of graph-based reconstruction tools, and proposing data-driven diagnostics for pipeline selection.

Although the species studied here (a non-model) has a draft assembly (Lin et al., 2021), it remains of limited quality for transcript-level applications because it is a fragmented draft in which only ≈71.3% of annotated genes are recovered as single-copy, implying substantial duplication or fragmentation in the remaining gene set, and they do not directly resolve full-length isoform structure or alternative splicing events. A high-quality reference transcriptome generated by long-read Iso-Seq is therefore still required to compensate for these genome-assembly limitations and to provide an isoform-resolved catalogue for reliable transcript-level quantification and downstream differential expression analysis.

In this study, we document the complete Iso-Seq transcriptome assembly pipeline for two organs, flower and leaf of *Moricandia arvensis* (*Brassicaceae*). A species displays broad phenotypic plasticity in both flowers and leaves, as well as in its C3-C4 photosynthetic metabolism (e.g., Gómez et al., 2020). Our aim was to generate a high-quality, non-redundant reference transcriptome for each organ, suitable for downstream differential expression analysis. To this, we combined PacBio Iso-Seq clustering with independent assembly strategies per organ and selected the strategy that best balanced three partially conflicting criteria; (i) transcriptome completeness, assessed as BUSCO completeness while retaining biological content; (ii) unique mapping rate, ensuring that reads can be confidently assigned to single transcripts; and (iii), low redundancy, avoiding inflated transcript counts that obscure quantification. These criteria compete with one another because retaining more transcripts increases completeness but also increases redundancy and multi-mapping, whereas aggressive collapsing improves mapping specificity at the risk of losing genuine isoforms. The quality of the selected assemblies was then validated in two independent ways, by open reading frame (ORF) prediction using TransDecoder, and by comparing selected annotated genes with published homologous sequences from *Arabidopsis thaliana* through multiple sequence alignment. Our workflow provides a reproducible framework for generating a reliable reference transcriptome for non-model plant systems, with improved transcript completeness compared to de novo short-read assemblies.

## MATERIALS AND METHODS

### Samples

Long-read Iso-Seq data and short-read RNA-seq data were generated from two organs (flower and leaf) of *Moricandia arvensis* under two environmental condition simulating Mediterranean spring (day/night = 10/14 h, temperature = 20/10 °C) and summer (day/night = 16/8 h, temperature = 30/20 °C). These two conditions were chosen to capture the high phenotypic plasticity of this species, which produces two markedly different flower phenotypes, one in each season, together with pronounced seasonal changes in leaves and in its C_3_-C_4_ photosynthetic metabolism (Gómez et al., 2020, 2024). A reference transcriptome built from both conditions is therefore expected to represent the full transcriptional repertoire of each organ, rather than that of a single seasonal phenotype. All plants were derived from a single natural population (Mar-33, Road A-385 Km 14, Granada, Spain; 37° 8’ 24” N, 3° 43’ 54” W). For long-read Iso-Seq, RNA was extracted using RNeasy Plant Mini Kit (Qiagen) from 5 individual plants per condition per organ combination and quantified using total RNA (Quant-iT RNA HS Assay kit) and RIN values. For each condition, each organ combination, equal amounts of high-quality total RNA from the five biological replicates were pooled into a single Iso-Seq sample. Pooling was used to maximise the diversity of transcripts and isoforms captured per library; because the aim of the Iso-Seq data was transcript discovery for a reference transcriptome, rather than variant-or individual-level analysis, the loss of individual resolution does not compromise downstream applications. High-quality RNA samples were pooled, and cDNA synthesis was conducted using the Iso-Seq Express 2.0 kit, followed by library construction using the Kinnex Full-Length RNA Kit (Suppl. Fig. 1A). Library preparation and PacBio sequencing were performed by Novogene, on Sequel II/IIe systems using the HiFi Revio system with SMRT Cell 25M Tray, yielding 13.4M – 15.8M reads per condition × organ combination. Raw PacBio Revio HiFi reads have been deposited in the European Nucleotide Archive under study accession PRJEB121201.

For short-read RNA-seq, we used previously published data from Gómez et al. (2020), generated from five individuals per condition and organ from the same population, but distinct from the plants used for Iso-Seq. Briefly, total RNA was extracted using the RNeasy Plant Mini Kit (Qiagen), libraries were prepared with the TruSeq Stranded Total RNA LT Sample Prep Kit (Plant), and sequencing was performed on an Illumina NovaSeq 6000 platform (paired-end, 150 bp) to a minimum depth of 40 M reads per sample; full details are given in the original publication, where the raw data are deposited (PRJNA604514). These short-read data were used to benchmark the Iso-Seq-derived reference transcriptome and to quantify expression, and the new reference was compared against the Trinity-assembled short-read reference from the same study.

### Iso-seq processing

High-quality Iso-Seq CCS reads were generated using the PacBio preprocessing pipeline. The Iso-Seq pipeline (v4.2.0) was used to process and cluster the sequences (Suppl. Fig. 1B). The workflow comprised: (i) CCS generation from raw subreads, (ii) primer removal using Lima v2.12.0, (iii) refinement by poly(A) trimming and artificial concatemer removal using Iso-Seq refine to produce full-length non-chimeric (FLNC) reads, and (iv) clustering of full-length reads to obtain high-quality (HQ) transcript clusters using Iso-Seq cluster (https://isoseq.how/clustering/cli-workflow.html).

To construct a comprehensive reference transcriptome for each organ, FLNC reads from both conditions (spring and summer) were combined prior to clustering, producing a single HQ clustered FASTA file per organ. A length filter of ≥500 bp was applied to both organ datasets for all downstream analysis.

### Sequence Redundancy Reduction

To remove sequence redundancy arising from minor allelic variation, minor sequencing differences, UTR length variation, and closely related paralogs, the HQ clustered FASTA was subjected to re-clustering using CD-HIT-EST v4.8.1 (Li and Godzik, 2006). Identity thresholds of 95% and 98% were evaluated. For each cluster, a single representative transcript was retained. Preliminary evaluation of the IsoSeq HQ clustered output and CD-HIT-EST re-clustered files was performed using BUSCO (Benchmarking Universal Single-Copy Orthologs; Manni et al., 2021) completeness assessment and isoforms-per-gene analysis (explained in a later section).

Biologically, stringency is also important because pooling RNA from multiple individuals (here, five biological replicates per condition × organ combination) introduces allelic variation that contributes to the redundancy pool without representing distinct gene transcripts. In addition, truncated reads, minor sequencing differences, and UTR-length variation in the same locus may inflate cluster counts. To establish the optimal pipeline, all assembly strategies were first evaluated on the flower dataset, after which the selected versions were applied to the leaf.

### Flower reference

#### Standardization of Collapsing Methods

Preliminary analysis of the Iso-Seq data revealed extreme transcript redundancy, with individual gene clusters containing over 1,000 transcripts at 95% nucleotide identity (CD-HIT re-clustered) (Suppl Fig.2). This level of redundancy may reflect a combination of real biological complexity (e.g., expanded gene families, allelic variation in a paleopolyploid genome (Lysak et al., 2005), alternative splicing), paralog co-clustering by CD-HIT, and fragmented or partial transcripts. To evaluate whether graph-based coding genome reconstruction could improve the reference, two pipeline versions were compared (Figure 1).

**Figure 1.**
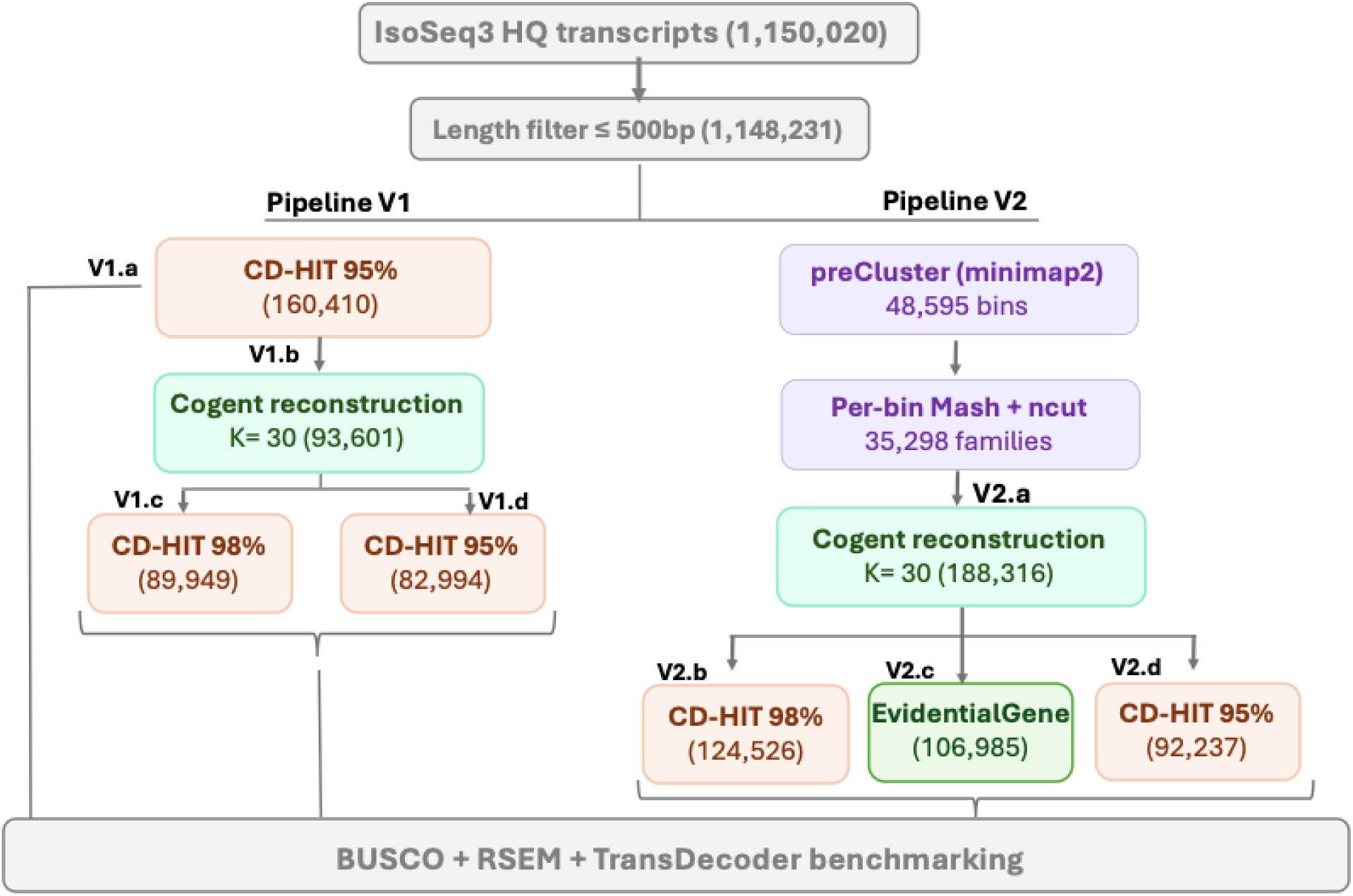
Workflow for Iso-Seq–derived full-length reference transcriptome construction and validation (In Flower organ). Schematic representation of the two pipeline versions evaluated in this study. Pipeline V1 (left) applies CD-HIT-EST identity-based clustering before Cogent graph reconstruction. Pipeline V2 (right) preserves all transcripts through a two-stage partitioning approach (minimap2 overlap binning followed by per-bin Mash clustering) before Cogent reconstruction. Both pipelines produce raw Cogent output that is subsequently processed through alternative redundancy reduction methods (CD-HIT at 95% and 98% identity, and EvidentialGene ORF-based classification). Numbers in parentheses indicate transcript counts at each stage; version sub-types are denoted by the lower-case letters as depicted. The curated reference transcriptome was used as a mapping target for short-read RNA-seq data to evaluate mapping efficiency.

##### Version 1 (V1)

CD-HIT pre-filter → Cogent (COding GENome Tool) reconstruction -- In V1, CD-HIT-EST at 95% identity was first applied to reduce the 1,148,231 length-filtered flower transcripts to 160,410 representative sequences, thereby reducing the extreme redundancy that would otherwise produce unmanageably large overlap graphs during Cogent reconstruction.

Genome-free transcript reconstruction was performed with Cogent v8.0.0 (https://github.com/Magdoll/Cogent/wiki), the 160,410 pre-filtered transcripts were directly subjected to Mash-based family finding using *run_mash.py* (k = 30, sketch size = 1,000, minimum distance = 0.95) to compute pairwise k-mer distances, followed by *process_kmer_to_graph.py* to apply normalised cut (ncut) graph partitioning into gene families. The “gene families” in Cogent’s terminology refer to transcript partitions inferred to derive from a single genomic locus, and do not necessarily correspond to gene families in the biological sense of evolutionarily related sets of paralogous genes. This direct Mash-based approach is appropriate for datasets of this size (≤ 200,000 transcripts) as recommended in the Cogent documentation.

Gene family consensus sequences were reconstructed using *reconstruct_contig.py* (default k-mer size = 30). Of the 27,218 partitions, 27,214 were successfully reconstructed (99.99%); families that initially failed due to graph cycle detection were recovered by re-running with *--nx_cycle_detection* at increasing kmer size (K = 40, 50, 80). Four partitions remained unresolved and were retained in the reference as their raw input sequences. The 27,214 reconstructed loci yielded 86,618 consensus transcripts. Combined 6,857 singleton transcripts and 126 raw transcripts from the four unresolved transcripts, the final V1 flower reference contained 93,601 transcripts. This raw Cogent output was then subjected to post-processing with CD-HIT-EST at 98% and 95% identity to evaluate the effect of further redundancy reduction (Figure 1).

##### Version 2 (V2)

Full dataset → preCluster → Cogent reconstruction --To assess whether the CD-HIT pre-filtering step in V1 might remove genuine biological variation before reconstruction, an alternative pipeline (V2) was evaluated that preserved all transcripts for partitioning. For the larger input dataset (1,148,231 transcripts), the Cogent documentation recommends a two-stage partitioning approach: first using minimap2 to roughly group transcripts into coarse bins, then applying the fine-grained Mash-based family finding within each bin.

A custom *run_preCluster.py* script was developed to perform minimap2 (v2.28) all-vs-all overlap mapping (preset *ava-pb*, minimum overlap: 200 bp, minimum identity: 0.85) on the 1,148,231 length-filtered transcripts, producing 48,595 coarse bins. Bin sizes ranged from 1 (singletons, n = 15,233) to 15,509 transcripts, with a median size of 4 transcripts.

Within each bin, fine-grained family finding was performed using Cogent’s *run_mash.py* with the same parameters as V1 (k = 30, sketch size = 1,000, minimum distance = 0.95). Processing was parallelised using GNU parallel (v20260122) with 30 concurrent jobs for bins ≤100 transcripts (n = 46,343), 10 concurrent jobs for bins of 101 transcripts (n = 2,243), and 4 concurrent jobs for bins >1,000 transcripts (n = 9). This stage produced 35,298 gene family partitions.

Gene family consensus sequences were reconstructed using *reconstruct_contig.py* (default k-mer size = 30), parallelised with GNU parallel (-j 60). Of the 35,298 partitions, 35,249 reconstructions succeeded on the first attempt. Of the 49 partitions that failed due to graph cycle detection, 46 were successfully recovered by re-running with *--nx_cycle_detection* and k = 40. Three partitions remained unresolvable: one family of 892 transcripts that failed at k = 220 after exhaustive internal retry, and two large partitions (5,099 and 8,262 transcripts) derived from a highly interconnected preCluster bin. For these three partitions, CD-HIT-EST at 99% identity was applied to collapse near-identical sequences, reducing them to 175, 1,057, and 2,923 representative transcripts, respectively. These sequences were included in the final reference as “unresolved” entries.

The combined V2 reference contained 188,316 transcripts: 184,161 Cogent-reconstructed consensus sequences from 35,295 gene families, plus 4,155 CD-HIT-99% representative sequences from the three unresolvable partitions.

#### Post-processing and reference selection

To evaluate the effect of post-processing stringency, the raw Cogent output from each pipeline version was subjected to redundancy reduction using CD-HIT-EST at 95% and 98% identity. Additionally, EvidentialGene tr2aacds4.pl (v2022.04.05; -NCPU 24, -MAXMEM 50000, -MINAA 30) (Gilbert, 2013) was applied to the V2 output to evaluate ORF-based transcript classification as an alternative to nucleotide-identity clustering.

V1 and V2 were evaluated as alternative strategies for balancing two opposing constraints: input scale (the number of transcripts requiring downstream partitioning, which grows with redundancy and directly determines Cogent’s computational and memory load) and graph-reconstruction tractability (whether the partitioned transcripts retain sufficient multi-isoform overlap per partition for Cogent’s De Bruijn graph algorithm to converge on a reliable consensus path). Aggressive pre-filtering reduces input scale but may erode tractability by removing the isoform diversity Cogent needs; minimal pre-filtering preserves diversity but exceeds Cogent’s computational limits. V1 applies aggressive identity-based pre-filtering using CD-HIT (95% sequence identity threshold) to reduce 1,148,231 length-filtered transcripts to 160,410 representative sequences before graph reconstruction. V2 retains all 1,148,231 transcripts and organises them using a two-stage partitioning approach: minimap2 ava-pb (for clustering by pairwise alignment) followed by per-bin Mash clustering (grouping sequences according to similarity metrics). Both pipelines were assessed by BUSCO completeness and RSEM (RNA-Seq by Expectation-Maximisation; Li & Dewey, 2011) mapping performance to determine which variant provides the most useful downstream reference (see Results).

### Leaf reference

The selected, V1 pipeline (CD-HIT 95% pre-filter → Cogent → CD-HIT) was evaluated on leaf tissue. CD-HIT-EST at 95% identity reduced the leaf IsoSeq HQ output to 130,407 representative transcripts. These were processed through Cogent v8.0.0 with the same parameters as for flower (Mash family finding at k = 30, sketch size = 1,000, minimum distance = 0.95; reconstruction with default k-mer size = 30), producing 65,567 consensus transcripts across 22,450 transcript partitions (gene families) (22,448 successfully reconstructed; 2 families included as unresolved raw transcripts).

### Quality Assessment

#### BUSCO Completeness Assessment

Transcriptome completeness was assessed using BUSCO v5.8.3 (Manni et al., 2021) in transcriptome mode with the brassicales_odb10 lineage dataset (n = 4,596 universal single-copy orthologues). Complete (C), single-copy (S), duplicated (D), fragmented (F), and missing (M) percentages were recorded. BUSCO assessment was performed at every processing stage to track the effect of each pipeline component on transcriptome completeness.

#### Short-Read RNA-seq mapping and Reference Validation

Illumina RNA-seq reads from all experimental conditions were quality-checked using standard filtering criteria to remove low-quality bases and adapter contamination, as described in Gómez et al. (2020). Cleaned reads were mapped to each Iso-Seq-derived reference variant using RSEM (v1.3.1) with Bowtie2 (Langmead & Salzberg, 2012) as the aligner, in paired-end mode. The Iso-Seq transcriptome reference was prepared using *rsem-prepare-reference* (https://deweylab.github.io/RSEM/rsem-prepare-reference.html) prior to quantification. Overall concordant alignment rate, unique mapping rate, and multi-mapping rate were recorded for each reference variant. This strategy enables transcript-level expression quantification in the absence of a reference genome, with RSEM’s expectation-maximisation algorithm handling multi-mapped reads probabilistically. A Trinity v2.8.4 (Grabherr et al., 2011) de novo assembly (424,981 transcripts; Gómez et al., 2020) was included as a short-read baseline for comparison.

#### Open Reading Frame Prediction

ORFs were predicted on both forward and reverse strands using TransDecoder (v5.5.0; Haas et al., 2013) based on coding potential and start codon refinement without homology support. For each transcript, the longest ORF and its completeness status (complete, 5′-partial, 3′-partial, internal) were recorded. ORFs with predicted scores >100 was initially evaluated in one sample to assess the effect of stringent filtering on transcriptome completeness (Suppl. Figure 3).

#### Functional Annotation

Functional annotation of the final selected reference transcriptome was performed using Trinotate (v 4.0.2; Bryant et al., 2017), incorporating homology search against SwissProt and Pfam domain identification.

#### Gene-Level Validation via Multiple Sequence Alignment

A subset of well-characterised genes was selected for validation: UBQ10 and PP2A as housekeeping reference genes; PAL1 and CHS key genes involved in phenylpropanoid and flavonoid biosynthesis, respectively; CAB1 as a chlorophyll a/b-binding protein; and RbcS1A as the small subunit of RuBisCO. For each gene, the best-matching transcript from the Iso-Seq reference transcriptome and the Trinity-based assembly was identified using tBLASTn (BLAST v2.13, Camacho et al., 2009) with a minimum e-value threshold of 1×10⁻^10^, querying against *Arabidopsis thaliana* protein sequences retrieved from UniProt. Open reading frames were predicted from the retrieved transcripts using TransDecoder (v5.5.0; https://github.com/TransDecoder) with a minimum ORF length of 50 amino acids. Predicted protein sequences were aligned with their *Arabidopsis* orthologues using MAFFT v7.562 (Katoh & Standley, 2013) with parameters --auto --reorder and visualised in Geneious Prime v2005.0.3 (https://www.geneious.com) relative to the *Arabidopsis* reference sequence.

The pipeline version showing the highest BUSCO completeness, best RSEM mapping performance, and strongest biological validation was selected as the final Iso-Seq reference transcriptome for downstream differential expression analysis.

## RESULTS AND DISCUSSION

### Iso-Seq Transcriptome Construction and Clustering

High-quality full-length reads generated by Iso-Seq were processed with the PacBio Iso-Seq pipeline to obtain polished, non-redundant isoforms (Suppl Fig. 4). Initial clustering yielded full-length HQ isoforms in combined conditions of flower (F_C: 1,150,020) and leaf (L_C: 803,912) (Suppl. Fig. 5). These sequences served as the primary long-read reference, although their high count indicated substantial redundancy. After applying a ≥ 500 bp length filter, 1,148,231 transcripts in flower and 802,754 in leaf was retained as input for further clustering (Suppl. Table 3). Long-read IsoSeq sequencing, therefore, yielded a substantial set of high-quality transcripts for both tissues, which were further refined through clustering to reduce redundancy.

### Iso-Seq HQ output and CD-HIT pre-filter selection

The raw Iso-Seq clustered HQ output contained 1,150,020 transcripts with 96.3% BUSCO completeness, an excellent starting point; however, 94.0% of BUSCO genes were duplicated, confirming extreme redundancy. Prior to committing to graph-based reconstruction, both CD-HIT identity thresholds were evaluated on the flower dataset to determine the appropriate pre-filter stringency (Figure 2 & Suppl Table 1). A pre-filter that collapses too aggressively risks eliminating genuine biological diversity (paralogs, splice variants), whereas one that collapses too weakly leaves Cogent unable to converge.

**Figure 2.**
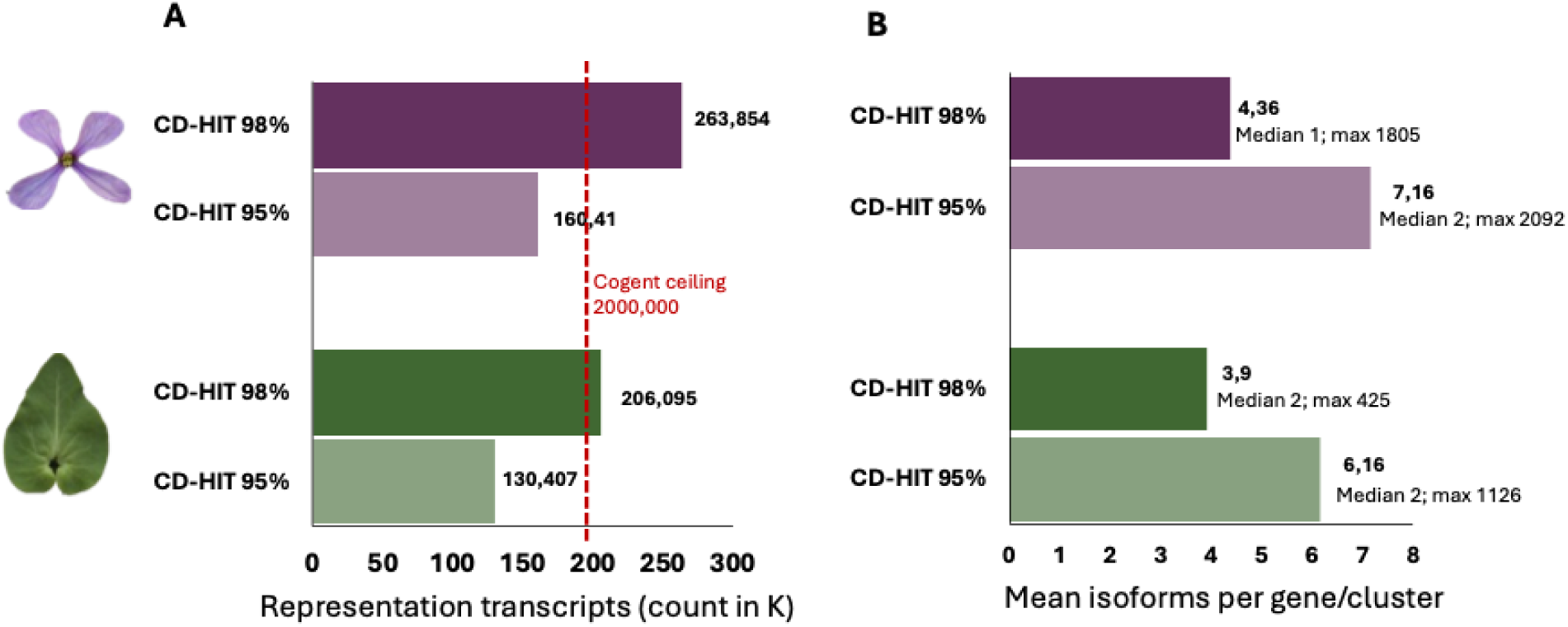
CD-HIT-EST identity-threshold evaluation for flower and leaf Iso-Seq HQ inputs. **(A)** Number of representative transcripts retained at each identity threshold, starting from 1,148,231 length-filtered Iso-Seq HQ transcripts. The red dashed line marks the ≤ 200,000-transcript ceiling for Cogent’s direct Mash-based family finding (run_mash.py); 98% output exceeds this ceiling for both tissues, ruling it out without an additional pre-processing step. **(B)** Mean isoforms per gene/cluster at each threshold, with median and maximum cluster size annotated where computed. The 95% threshold preserves the multi-isoform evidence required for reliable Cogent graph reconstruction (median cluster size ≥ 2 in flower); the 98% threshold collapses most gene families to single representatives (median = 1 in flower).

CD-HIT at 98% identity reduced the 1,148,231 length-filtered transcripts to 263,854 representatives (flower) and 206,095 (leaf), with mean isoforms per gene of 4.36 and 3.90, respectively, and a maximum cluster size of 1,805 (flower). CD-HIT at 95% identity produced 160,410 flower and 130,407 leaf representatives, with mean isoforms per gene of 7.16 and 6.16, and maximum cluster sizes of 2,092 and 1,126, respectively (Figure 2A & B). Two criteria determined the selection of the 95% threshold for downstream Cogent input. First, the Cogent documentation specifies that direct Mash-based family finding (run_mash.py) is appropriate for datasets of ≤200,000 transcripts; the 98% output of 263,854 sequences exceeds this ceiling, rendering it incompatible with direct Mash partitioning without an additional pre-processing step. Second, the 98% threshold retained a median cluster size of 1 in flower, indicating that most gene families were represented by a single transcript and therefore lacked the multi-isoform evidence required for reliable Cogent graph reconstruction. The 95% threshold, yielding 160,410 sequences within the tractable range and a median cluster size of 2, was therefore selected as the Cogent pre-filter for flower tissue. The same rationale applied to leaf, where the 95% output (130,407 sequences) was within the Cogent-compatible range, and the 98% output (206,095) was not.

### Flower Transcriptome

The downstream pipeline must contend with two intertwined sources of complexity that the pre-filter alone cannot disentangle. Biologically, the flower transcriptome legitimately carries multiple isoforms per gene, expanded paralog families from ancestral duplications, and allelic variation inflated by our five-individual pooling. Technically, CD-HIT collapses by sequence identity without distinguishing these sources, and Cogent’s graph reconstruction requires that genuine multi-isoform support survives the collapse. The following analysis evaluates how well the flower assembly navigates this trade-off.

#### Two clustering-reconstruction strategies: V1 and V2

V1 (CD-HIT 95% pre-filter → Cogent → CD-HIT): Here we used 160,410 reduced transcripts using 95% identity clustering, then applied Cogent reconstruction on the pre-filtered sequences, then applied a further CD-HIT step to the Cogent output (Figure 1).

The version V1 generated reference transcriptome (all versions V1.b to V1.d) has preserved and retained BUSCO completeness (95%) comparable to the un-collapsed set (Iso-Seq HQ) and showed balanced read-mapping performance (77.7 to 77.5%), indicating that the pre-filtering and graph reconstruction workflow was appropriate for the flower dataset (Figure 3A & B, Table 1). Notably, V1.b has lower total mapping (77.7 ± 2.4%) than CD-HIT 95% (82.7 ± 3.3%), which is expected and not a quality concern. A more tightly collapsed assembly maps fewer reads in total but maps them far more confidently. Moreover, we are working with tissue/organ-specific transcriptomes, so a high percentage of alignment is not expected.

**Figure 3.**
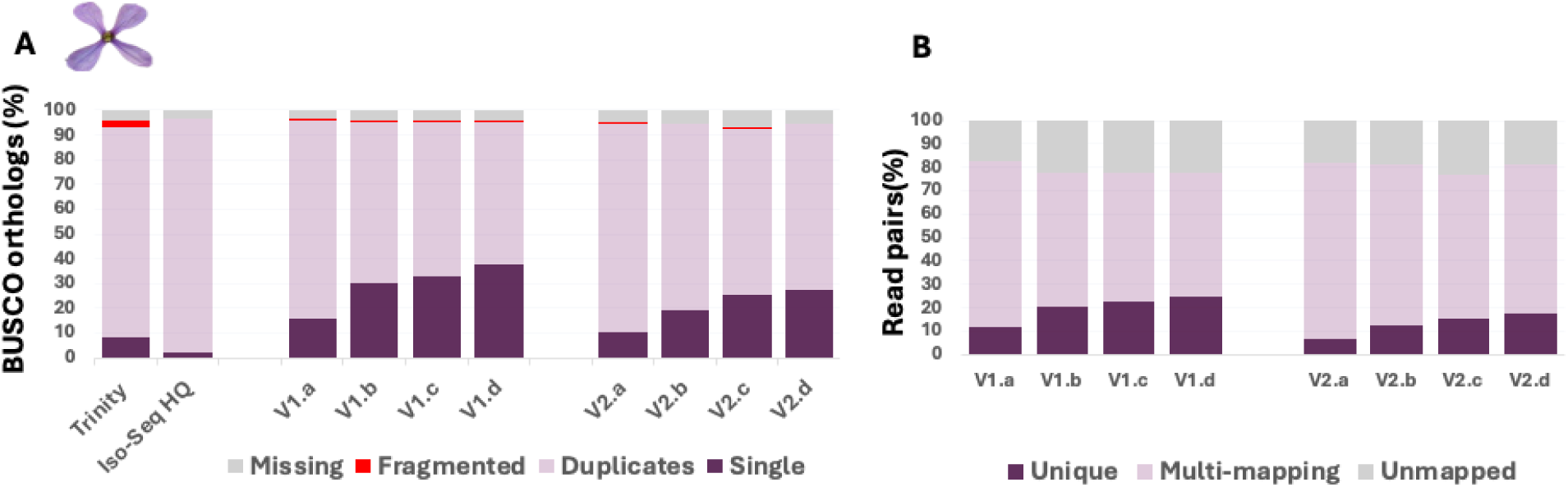
Quality assessment of flower transcriptome assembly variants. **(A)** BUSCO completeness against the *brassicales_odb10* lineage dataset (n = 4,596 orthologues) for all V1 and V2 pipeline variants. Stacked bars show the proportion of single-copy (S), duplicated (D), fragmented (F) and missing (M) BUSCO orthologues. **(B)** RSEM/Bowtie 2 concordant alignment rates for paired-end RNA-seq reads, shown as the mean of two biological replicates (one spring, one summer individual) mapped to each leaf V1 variant. Bar height represents total mapping rate; the dark portion within each bar represents the unique mapping fraction. The Trinity short-read assembly is included as a baseline.

**Table 1.** Iso-Seq transcriptome assembly metrics for flower pipeline variants. Values shown for each V1 and V2 variant include transcript count, BUSCO completeness (single-copy, duplicated, fragmented, missing) against *brassicales_odb10* (n = 4,596), and RSEM/Bowtie 2 total and unique mapping rates against the matching short-read RNA-seq library (average of two replicates ± SD).

| Assembly stage | Transcripts | BUSCO C% | BUSCO S% | BUSCO D% | Total map% | Unique map% |
| --- | --- | --- | --- | --- | --- | --- |
| IsoSeq HQ | 1,150,020 | 96.3 | 2.3 | 94.0 | — | — |
| V1.a | 160,410 | 95.9 | 15.6 | 80.3 | $82.7 \pm 3.3$ | $11.9 \pm 1.7$ |
| V1.b | 93,601 | 95.3 | 29.9 | 65.4 | $77.7 \pm 2.4$ | $20.3 \pm 1.3$ |
| V1.c | 89,949 | 95.3 | 32.9 | 62.4 | $77.7 \pm 2.4$ | $22.4 \pm 1.5$ |
| V1.d | 82,994 | 95.3 | 37.8 | 57.5 | $77.5 \pm 2.4$ | $24.5 \pm 1.9$ |
| V2.a | 188,316 | 94.3 | 10.3 | 84.0 | $81.6 \pm 2.0$ | $7.0 \pm 0.5$ |
| V2.b | 124,526 | 94.3 | 19.1 | 75.2 | $81.4 \pm 2.0$ | $12.5 \pm 1.1$ |
| V2.c | 106,985 | 92.7 | 25.3 | 67.4 | $77.0 \pm 0.3$ | $15.4 \pm 0.9$ |
| V2.d | 92,237 | 94.2 | 27.3 | 66.9 | $80.9 \pm 2.1$ | $17.7 \pm 1.7$ |

V1.a, Iso-Seq re-clustered with CD-HIT 95%; V1.b, Cogent-reconstructed from V1.a; V1.c, V1.b post-processed with CD-HIT 98%; V1.d, V1.b post-processed with CD-HIT 95%; V2.a, Cogent-reconstructed from full Iso-Seq HQ input; V2.b, V2.a post-processed with CD-HIT 98%; V2.c, V2.a classified by EvidentialGene tr2aacds4.pl; V2.d, V2.a post-processed with CD-HIT 95%.

Further, to evaluate the effect of post-processing stringency, CD-HIT-EST was applied to the raw Cogent output at two identity thresholds, V1.c (CD-HIT 98%) and V1.d (CD-HIT 95%). Interestingly, both maintained the BUSCO completeness 95.3% like V1.b, although single copy percentage increased with a rise in threshold level and vice-versa with duplicates in BUSCO (Figure 3A and Table 1). Moreover, the unique mapping rate is increased with CD-HIT percentage threshold level, 22.4 ± 1.5% at CD-HIT 98% and 24.5 ± 1.9% at CD-HIT 95% (Figure 3B & Table 1). The highest yield in unique mapping rate came at no gain in total mapping, consistent with over-collapse of paralogs at the 95% post-processing step.

*V2 (IsoSeq directly → Cogent → CD-HIT & EvidentialGene):* To assess whether the CD-HIT pre-filtering step might remove genuine biological variation before reconstruction, we also evaluated an alternative pipeline (V2) that preserved all transcripts through a two-stage partitioning approach (Figure 1). All 1,148,231 length-filtered transcripts were first subjected to minimap2 all-vs-all overlap alignment, producing coarse bins based on shared sequence (48,595 bins; parameters: -x ava-pb, minimum overlap 200 bp, minimum identity 85%). Within each bin, Cogent’s Mash-based k-mer distance estimation and normalised cut graph partitioning identified 35,298 transcript variants (gene family partitions). Cogent reconstruction of these partitions produced 188,316 consensus transcripts from 35,295 transcript variants (gene families) (V2.a). This V2 raw output was subjected to the same post-processing variants evaluated for V1.

The V2 pipeline achieved higher total mapping rates (80–81%) compared to V1 (77.7%), suggesting that CD-HIT pre-filtering in V1 removed some genuine transcript sequences (Figure 3B). However, V1 consistently outperformed V2 on BUSCO completeness (95.3% vs 94.2–94.3%) (Figure 3A & Table 1) and unique mapping (20–24.5%; V1.b, V1.c, V1.d vs 12–17.7%; V2.b, V2.d), indicating that the simpler graph structures produced by pre-filtered input led to higher-quality consensus sequences for expression quantification (Figure 3B & Table 1). The reason may be structural: V2 fed 7 times more sequences into Cogent, creating noisier gene family bins. Cogent’s reconstruction within each bin was less accurate because bins contained more bogus members, producing more redundant output. The CD-HIT 95% pre-filter in V1 was essential preparation, as it removed trivial duplicates so Cogent could focus on genuine isoform diversity within each family.

Notably, EvidentialGene’s ORF-based classification (tr2aacds4.plv2022.04.05), while designed for short-read assemblies, reduced BUSCO completeness to 92.7% when applied to Iso-Seq data (V2.c), indicating that its protein quality filters are overly aggressive for full-length long-read transcripts. Both Cogent graph reconstruction (35,295 gene families) and EvidentialGene ORF classification (35,229 gene loci) independently converged on ≈ 35,200 gene loci, providing confidence in the estimated gene count for paleopolyploid *M. arvensis* (Lysak et al., 2005).

V1.c (Cogent + CD-HIT 98%) was retained as the final flower reference. Compared to V1.b (Cogent raw), V1.c maintained equivalent BUSCO completeness (95.3%) but with reduced duplication and elevated single-copy proportion (Figure 3A, Table 1), indicating effective removal of near-identical paralogs without loss of biological content captured during graph reconstruction. Compared to V1.d (Cogent + CD-HIT 95%), V1.c retained similar total RSEM alignment compared to Cogent raw, which is the relevant metric for downstream RSEM-based quantification: the expectation-maximisation algorithm probabilistically assigns multi-mapped reads to specific transcripts, making total mapping the primary determinant of expression quantification accuracy. V1.c therefore offers the best balance of completeness, redundancy reduction, and mapping performance among the variants evaluated.

A short-read Trinity assembly (424,981 transcripts, BUSCO 93.2% complete) was also evaluated as a baseline comparison. While Trinity achieved the highest total mapping rate (87.09%), its extreme redundancy (85.4% duplicated BUSCOs) and low unique mapping (13.39%) render it unsuitable as a reference for expression quantification without substantial post-processing compared with Iso-Seq.

### Leaf Transcriptome

#### Evaluation of Cogent reconstruction for leaf tissue

After thorough evaluation of all versions in flower organ transcripts, the same V1 pipeline (CD-HIT 95% pre-filter to Cogent reconstruction) was applied to leaf tissue. The raw IsoSeq clustered HQ output for leaf contained 803,912 transcripts with 90.8% BUSCO completeness. CD-HIT 95% reduced the leaf IsoSeq output to 130,407 transcripts (V1.a), and Cogent reconstruction produced 65,688 consensus transcripts (V1.b) across 22,450 transcript variants (gene families) (Table 2). The IsoSeq HQ transcripts showed 90.8 % BUSCO completeness, which is less than that of the flower organ, whereas CD-HIT 95% again preserved the BUSCO completeness to 90.1% with enhanced single-copy genes from 4.1% to 20.7% (Figure 4A & Table 2). However, BUSCO assessment revealed a substantial loss of transcriptome completeness during Cogent reconstruction (V1.b). It reduced BUSCO completeness from 90.1% to 78.0%, while missing genes tripled from 9.0% to 21.2% (976 conserved orthologues) in V1.b (Figure 4A & Table 2). However, single-copy gain is also not substantial. Post-processing with CD-HIT 98% (V1.c) did not contribute to this loss, confirming that the gene loss occurred during the graph reconstruction step.

**Figure 4.**
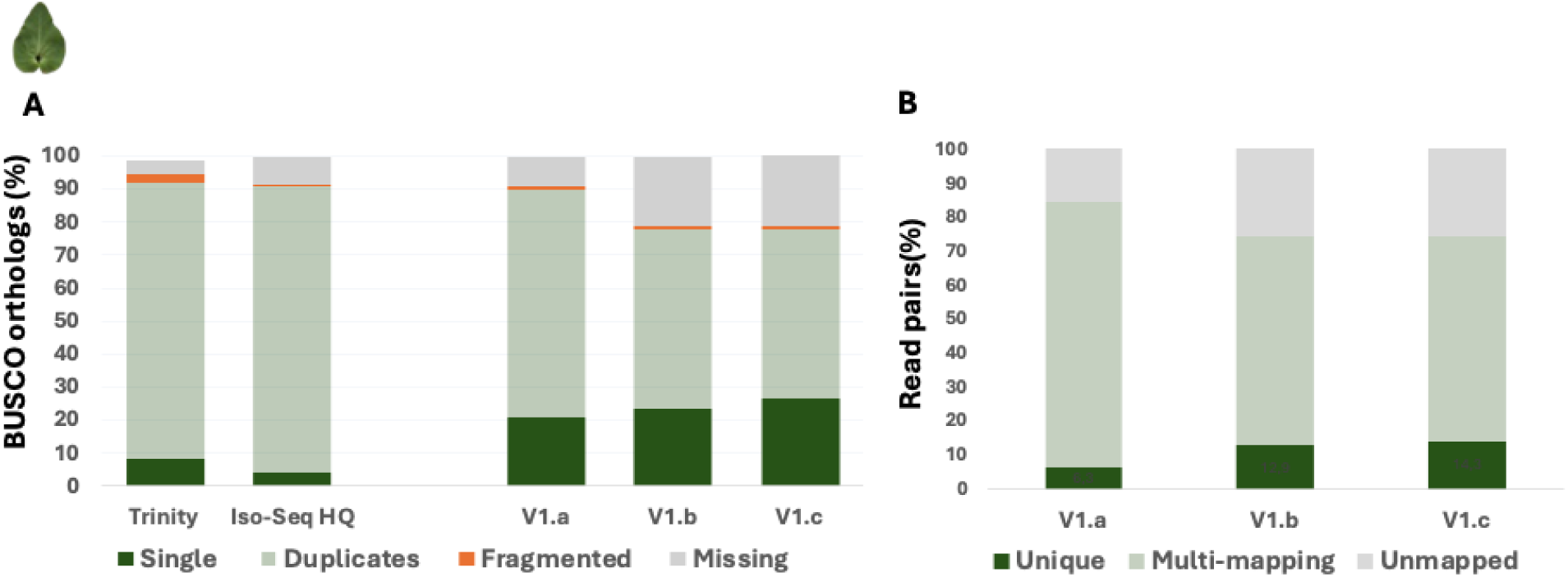
Quality assessment of leaf transcriptome assembly variants for the V1 pipeline. **(A)** BUSCO completeness against the *brassicales_odb10* lineage dataset (n = 4,596 orthologues). Stacked bars show the proportion of single-copy (S), duplicated (D), fragmented (F) and missing (M) BUSCO orthologues across V1 variants and the Trinity short-read baseline. **(B)** RSEM/Bowtie 2 concordant alignment rates for paired-end RNA-seq reads, shown as the mean of two biological replicates (one spring, one summer individual) mapped to each V1 variant. Bar height represents total mapping rate; the dark portion within each bar represents the unique mapping fraction.

**Table 2.**
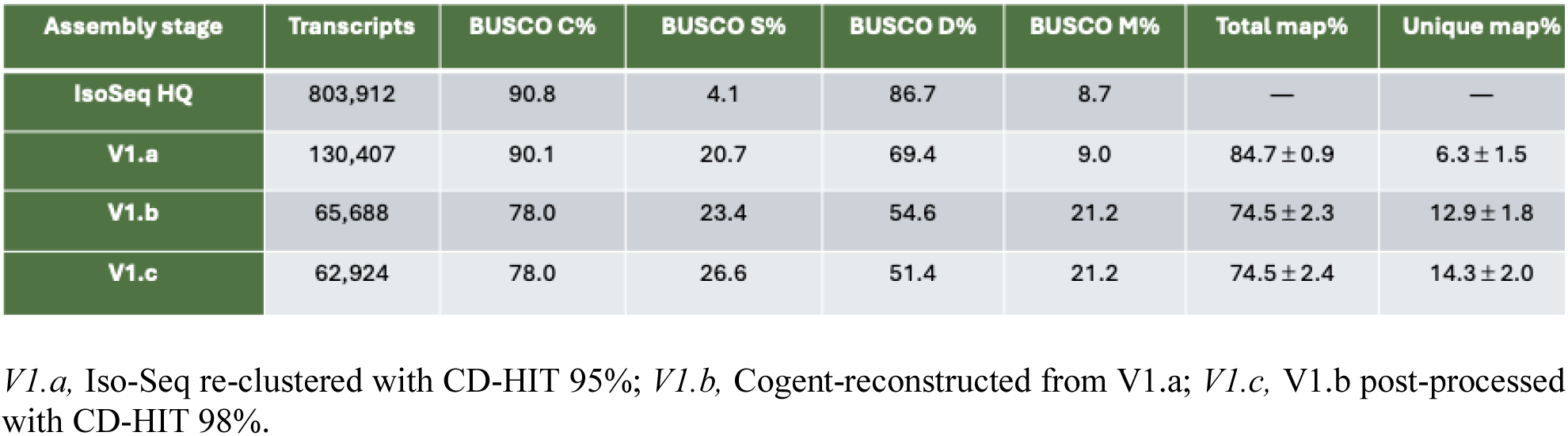
Iso-Seq transcriptome assembly metrics for leaf V1 pipeline variants. Values shown for each variant include transcript count, BUSCO completeness against *brassicales_odb10* (n = 4,596), and RSEM/Bowtie 2 total and unique mapping rates (average of two replicates).

Concurrently, the unique mapping gain was only 6.6 percentage points (6.3 ± 1.5% vs 12.9 ± 1.8%), while total mapping rate fell by 10.2 percentage points from 84.7 ± 0.9% to 74.5 ± 2.3% (Figure 4B & Table 2), indicating that real biological content was lost during Cogent reconstruction in the leaf dataset.

#### Causes for Cogent failure in leaf organ

We identified two main factors responsible for this failure: (i) incorrect gene merging and (ii) extreme expression skew.

First, analysis of the isoform distribution provided mechanistic insight into the suboptimal performance of Cogent in leaf tissues (Figure 4 & Suppl Table 1). The key diagnostic metric is the median isoform count, which increased from 2 to 4 after Cogent processing, a trend opposite to expected behaviour (Figure 5D). In an optimal functioning (as seen in flower), Cogent preserves the overall distribution shape while rescaling it downward by proportionally reducing singletons, maintaining the median isoform count, and collapsing large gene clusters (Figure 5A, B). In the leaf data, the distribution profile was inverted. Singleton genes decreased sharply from 58,331 (44.7%) to 5,987 (9.1%), representing a net loss of 52,344 genes (Supplementary Table 2). Concurrently, low-isoform bins expanded (Figure 5D); for example, the 3-isoform bin increased from 10,463 to 10,590, and the 4-isoform bin grew from 6,694 to 8,008 (Figure 5C). This pattern is diagnostic of incorrect gene merging.

**Figure 5.**
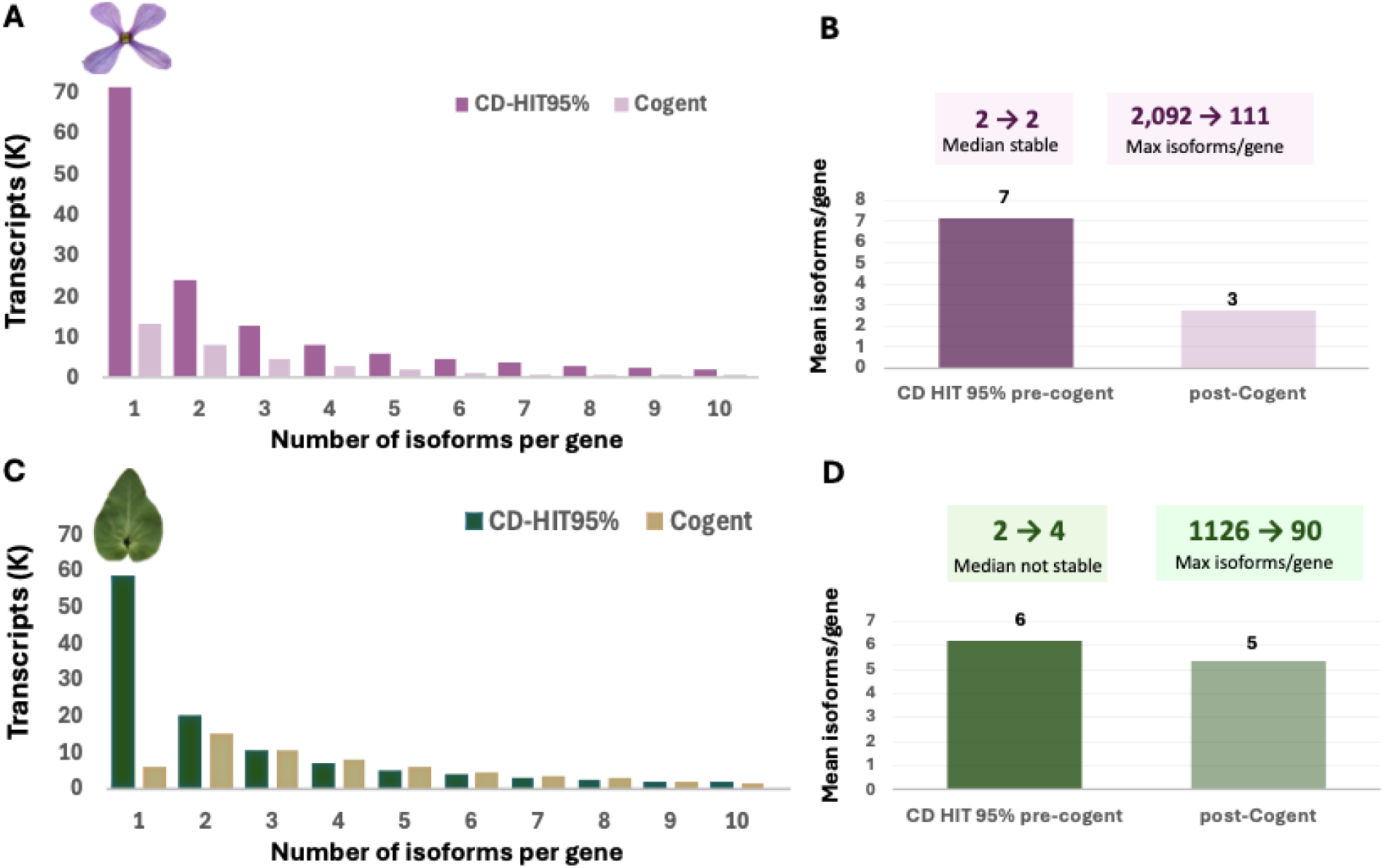
Per-gene isoform distribution before and after Cogent reconstruction. (**A, C**). Number of isoforms per gene (Cogent-reconstructed) or per cluster (CD-HIT 95% clustered) in flower (A) and leaf (C); each panel compares the CD-HIT 95% clustered transcriptome with the Cogent-reconstructed transcriptome. (**B, D**) Mean, median and maximum isoforms per gene/cluster from the CD-HIT clustered to the Cogent-reconstructed output for flower (B) and leaf (D).In the leaf data, where 44.7% of genes have only one transcript, these singletons were being pulled into bins with their nearest neighbors, genuinely distinct genes that happen to share some sequence similarity (Suppl Table 2 & 3). Cogent then reconstructed these mixed bins as multi-isoform families, splitting and merging transcripts incorrectly. The result was fragmentation of real genes (explaining the BUSCO completeness loss) and artificial inflation of isoform counts (explaining the median increase).

Second, the distribution of full-length (FL) reads across transcripts pointed to the same problem. Although both tissues were sequenced to comparable depths (∼28 million FL reads) and shared an identical median support of 4 FL reads per transcript (Table 3, Figure 6A), their underlying distribution profiles diverged in a critical manner. Despite identical median depth, leaf’s top 1% of transcripts consumed 59.8% of all FL reads compared to 49.2% in flower (Table 3 & Figure 6B). This means the remaining 99% of leaf transcripts competed for only 40.2% of reads, leaving most genes with insufficient multi-isoform coverage for Cogent’s graph reconstruction. The cause is biological; the leaf had 236 hyper-expressed transcripts (max 272,794 FL reads) versus the flower’s 108 (max 63,830), indicating extreme expression skew driven by organ-specific highly abundant transcripts such as photosynthesis-related genes in the leaf organ (Table 3). Additionally, the lower completeness compared to flower (96.3%) and the higher BUSCO missing rate (8.7% vs ≈3.7% for flower) already indicated that the leaf had lower per-gene sequencing depth.

**Figure 6.**
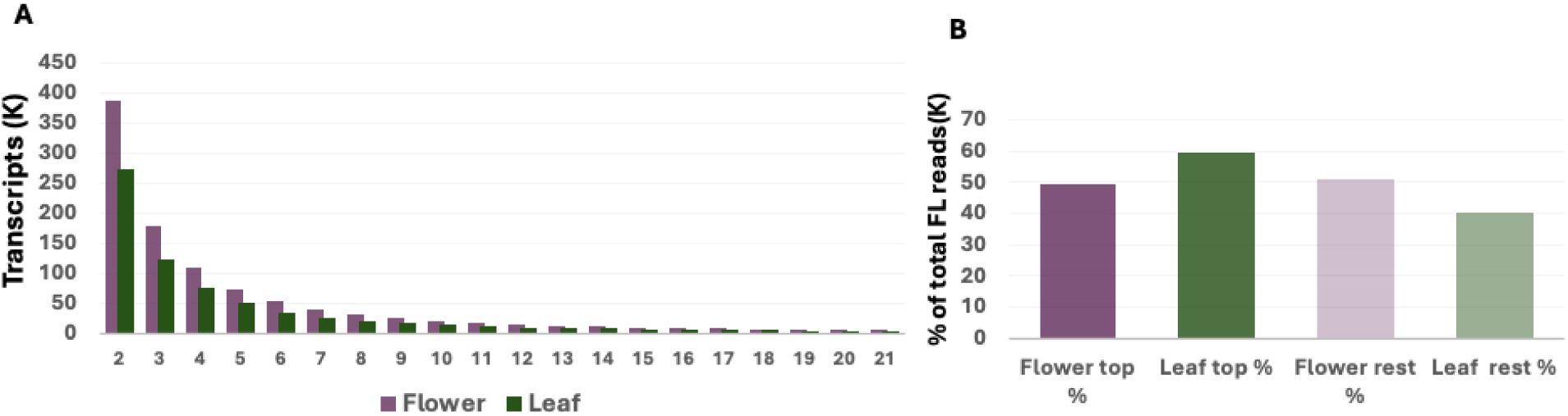
Full-length (FL) read distribution and expression skew in flower and leaf Iso-Seq data. (**A**) Distribution of FL-read support per transcript across the 1–20 FL-read range, showing the per-transcript depth profile in each tissue. (**B**) Cumulative FL-read share captured by the top-ranked transcripts in each tissue, illustrating the disproportionate share consumed by leaf’s most highly expressed transcripts.

**Table 3.** Full-length (FL) read distribution metrics for flower and leaf Iso-Seq data. Values shown include total FL reads, median FL reads per transcript, maximum FL reads per transcript, number of hyper-expressed transcripts, and the FL-read share captured by the top 1% of transcripts in each tissue.

| Metric | Flower | Leaf | Interpretation |
| --- | --- | --- | --- |
| Total FL reads | 28,536,006 | 28,085,422 | Comparable depth |
| Median FL/transcript | 4 | 4 | Identical |
| Max FL reads | 63,830 | 272,794 | Leaf 4.3x more skewed |
| Transcripts >10k FL reads | 108 | 236 | 2.2x more in leaf |
| Top 1% transcripts — FL share | 49.2% | 59.8% | Leaf more concentrated |
| Remaining 99% — FL share | 50.8% | 40.2% | Less depth for most genes |

Running V2 on leaf (feeding more sequences into Cogent without pre-filtering) would worsen the problem, not improve it. V2 demonstrated inferior performance even for flower tissue, where Cogent worked correctly. For leaf, feeding 803,912 unfiltered sequences into Cogent would produce even noisier bins, more incorrect merges, and greater BUSCO loss. Thus, this path was not pursued.

#### Selection of leaf reference

Given that Cogent reconstruction was demonstrably detrimental to leaf transcriptome quality and V2 was not viable for the same input characteristics, the leaf reference was selected from the available CD-HIT pre-filtered output without graph reconstruction. The CD-HIT 95% direct output (130,407 transcripts, designated V1.a) preserved BUSCO completeness at 90.1%, which is comparable to the unprocessed Iso-Seq HQ input and substantially higher than the post-Cogent V1.b reference at 78.0%. RSEM mapping showed 84.7 ± 0.9 % total alignment rate, with 6.3 ± 1.5% unique mapping. Although unique mapping is lower than the flower V1.c reference (22.4 ± 1.5%), this reflects the absence of graph-based isoform collapse rather than a quality deficit; for RSEM-based downstream analysis, the expectation-maximisation algorithm probabilistically resolves multi-mapped reads, making total mapping rate the primary determinant of expression quantification accuracy. The V1.a reference, therefore, retains its biological content for differential expression analysis at the cost of higher per-transcript mapping ambiguity and was retained as the leaf sequence reference for the present study. However, unlike the flower V1.c reference, in which Cogent reconstruction yields transcripts already organised into gene-level families with isoforms, the leaf V1.a reference is a regular, non-redundant set of representative transcripts without gene-level grouping. Because downstream functional annotation and gene-level differential expression require transcripts to be grouped into genes and their constituent isoforms, a separate gene-clustering step was applied to the leaf reference (Suppl. Material 1).

#### Read-clustering recovers leaf gene-isoform structure

Read-clustering recovers leaf gene-isoform structure where graph reconstruction failed. Because Cogent reconstruction degraded the leaf transcriptome (as explained above), the leaf V1.a reference lacked the gene-level organisation that Cogent provided for flower. Two read-clustering strategies were therefore compared on the identical 130,407-transcript V1.a set (Suppl. Table 4).

Expression-aware clustering with short-read RNA-seq (Corset; Davidson & Oshlack, 2014) grouped 121,235 transcripts into 43,751 gene clusters, with 44.2% singletons and a smooth size distribution (largest cluster, 90 transcripts). The representative set (one transcript per cluster) retained 86.4% BUSCO completeness (single-copy 48.7%, duplicated 37.6%, missing 11.5%), close to the 90.1% of the full V1.a set and far above the 78.0% of the Cogent-reconstructed V1.b, indicating that clustering preserved distinct conserved genes rather than merging them. This expression-aware clustering (short-read Corset) preserved both completeness and retained the duplicated-gene structure of a paleopolyploid genome, separating paralogues based on their divergent expression across spring and summer.

Meanwhile, sequence-based clustering (isONclust3; Petri and Sahlin, 2025) grouped all 130,407 transcripts into 37,344 clusters (39.5% singletons; largest cluster, 134 transcripts), with 83.8% representative BUSCO completeness. Although completeness was comparable to Corset, the single-copy and duplicated proportions differed markedly: isONclust3 returned 70.1% single-copy and only 13.7% duplicated BUSCOs, against 48.7% and 37.6% for Corset. For a paleopolyploid genome (Lysak et al., 2005), in which conserved orthologues are expected to be present in multiple paralogous copies, the substantially lower duplication of isONclust3 is consistent with sequence similarity alone merging paralogous copies into single clusters, collapsing genuine duplicates that the expression-aware method kept separate. A concrete example is the UBQ10 family: two divergent family members (transcript/422248 and transcript/567399) were placed in separate Corset clusters but merged into a single isONclust3 cluster.

The short-read Corset clustering was therefore selected as the leaf gene model. This interpretation rests on the paleopolyploidy expectation that paralogous copies should be retained as separate genes and on a single illustrative family (UBQ10); the lower isONclust3 duplication could alternatively reflect legitimate collapse of true redundancy. Corset assigned 121,235 of the 130,407 transcripts to gene clusters; the ∼9,000 transcripts below the minimum-count threshold (-m 10) were retained in the reference as unassigned singletons. Transcripts corresponding to the genes used for sequence-level validation (PP2A, PAL1, CHS, CAB1, RbcS1A) each resolved into small clusters of one to five transcripts, consistent with correct gene-boundary assignment. UBQ10 was represented through three family members distributed across three small clusters; its two most repeat-identical copies (transcript/570410, transcript/118010) received no uniquely assignable short reads and were absent from the clustering, reflecting the established difficulty of short-read quantification across the polyubiquitin tandem-repeat array, the same structural feature that long-read Iso-Seq resolves unambiguously.

### Full Length Completeness

An independent validation of pipeline quality comes from TransDecoder completeness analysis, which classifies each transcript as complete (both 5’ and 3’ ends present), 5’-partial, 3’-partial, or internal (both ends missing). This analysis confirms the pipeline decisions independently of BUSCO or mapping statistics.

In flowers, the improvement is monotonic and striking; complete transcripts rise from 42.0% (Trinity) through 57.1% (Iso-Seq HQ) and 80.5% (CD-HIT NR) to a peak of 85.2% at Cogent (Figure 7A). Internal fragment rate collapses from 19.5% (Trinity) to 0.1% (Cogent). The leaf data show the same monotonic improvement profile: complete transcripts rise from 40.7% (Trinity) through 57.7% (Iso-Seq HQ) and 81.3% (CD-HIT NR) to 86.9% at the Cogent stage (Figure 7B). At the leaf Cogent stage, the proportion of complete ORFs was high (86.9%), confirming that Cogent successfully reconstructed full-length coding sequences from the input transcripts.

**Figure 7.**
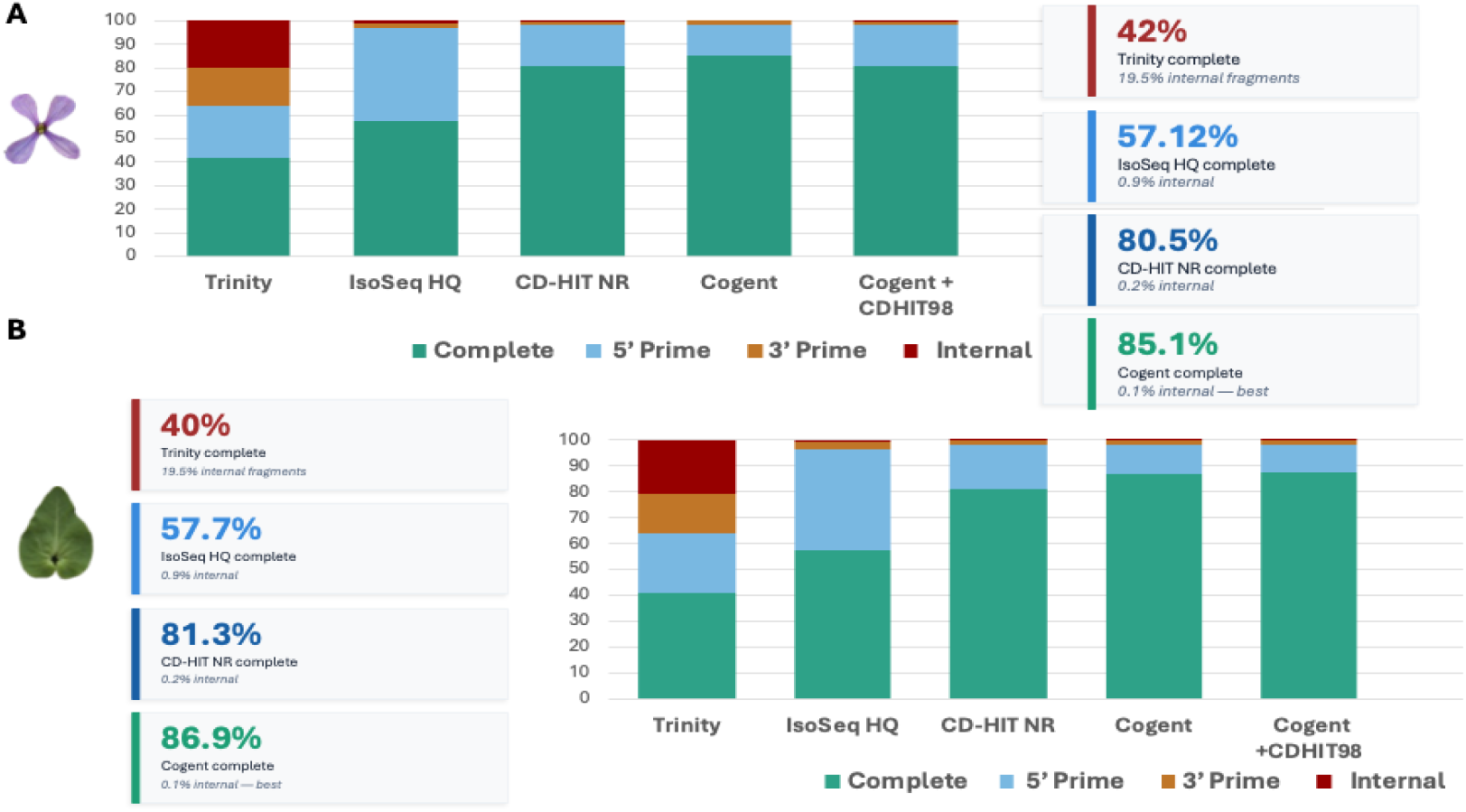
TransDecoder-predicted coding-potential profile across pipeline stages. Proportion of transcripts classified as complete (both 5′ and 3′ ends present), 5′-partial (3′ end missing), 3′-partial (5′ end missing), and internal (both ends missing) for each processing stage, Trinity short-read assembly, Iso-Seq HQ clustered, CD-HIT 95% non-redundant (CD-HIT NR) and Cogent-reconstructed in (A) flower and (B) leaf. were preferentially the longer, full-length representatives of each gene family, exactly the behaviour expected from an Iso-Seq-based assembly. In short, the leaf problem is one of gene grouping, not of ORF reconstruction.

This finding is informative because it locates the source of the leaf BUSCO loss. The loss did not arise from Cogent failing to recover ORF structure within each reconstructed gene family; rather, it arose from distinct genes being incorrectly grouped into the same family before reconstruction. The TransDecoder result thus complements the BUSCO evidence: at every pipeline stage, and in both tissues, the transcripts retained

### Validation Using Known Gene Sequences

To independently validate the biological accuracy of the Iso-Seq reference transcriptome, predicted protein sequences from six target genes were aligned against their *Arabidopsis thaliana* orthologues and compared with Trinity-assembled equivalents (Suppl. Fig. 8). Iso-Seq-derived transcripts consistently recovered proteins of expected length and high sequence identity across both tissues. CAB1, PAL1, CHS and RbcS1A Iso-Seq sequences aligned with *Arabidopsis* orthologues across their full length with minimal gaps, confirming complete transcript capture. UBQ10 from the leaf Iso-Seq reference achieved 100% amino acid identity to the *Arabidopsis* orthologue, the highest identity observed across all genes examined. The flower Iso-Seq UBQ10 transcript (685 aa) represents the full-length polyubiquitin precursor, a tandem repeat structure correctly assembled by Iso-Seq long reads that would be difficult to reconstruct from short-read data. Two exceptions within the Iso-Seq flower reference warranted further attention. PP2A recovered only a partial protein (185 aa vs expected ≈ 305 aa) from the flower reference, most likely reflecting low per-gene FL-read coverage for this regulatory phosphatase in flower or can be a fragmented isoform case, consistent with PP2A’s characteristically low expression in flower organs relative to vegetative tissues (Csordás et al., 2000; Máthé et al., 2023). However, the leaf reference recovered a complete PP2A protein (337 aa, 100% query coverage), consistent with higher housekeeping gene expression in vegetative organs and confirming that the leaf V1.a reference accurately captures constitutively expressed genes.

Trinity-assembled transcripts showed marked organ-specific chimerism and truncation. CAB1 Trinity contigs were 455 aa (flower) and 541 aa (leaf) compared to the expected 267 aa, reflecting assembly of non-homologous sequences into chimeric transcripts, a well-documented artefact of short read de novo assembly in the presence of conserved domains. Trinity_LF_RbcS1A (593 aa) similarly exhibited severe chimerism. PAL1 from the Trinity flower assembly was truncated at 706 aa compared to 722 aa in the Iso-Seq reference and *Arabidopsis* orthologue (725 aa), consistent with the lower BLAST identity observed for this contig (80.8% vs 93.4% for Iso-Seq). By contrast, CHS (325 aa) showed equivalent structure across both reference types, showing that the Iso-Seq advantage is most pronounced for larger, more complex transcripts prone to mis-assembly.

Collectively, these results provide independent sequence-level validation of the Iso-Seq reference transcriptome and corroborate the pipeline benchmarking findings, Iso-Seq-derived references recover complete, accurately bounded transcripts with higher fidelity to known orthologues than Trinity-based assemblies, particularly for transcripts susceptible to chimeric joining or 5′/3′ truncation during short-read assembly. De novo transcriptome assembly from short reads (e.g., Trinity) has been recognised as having difficulties in recovering complete transcripts and correctly resolving isoforms due to ambiguities in the sequence graph and erroneous chimeric assemblies (Steijger et al., 2013).

### Pipeline performance is organ-specific

The same pipeline that produced the flower V1.c reference at 95.3% BUSCO and 22.4 ± 1.5% unique mapping reduced leaf BUSCO completeness from 90.1% to 78.0% (a loss of 976 conserved orthologues). This divergence within a single species is the central methodological finding of this study and demonstrates that long-read reference construction is not a one-size-fits-all workflow. The cause lies in two organ-specific characteristics of the input data. Median full-length read support per transcript was identical between tissues (4 reads in both), but the distributional shape differed: leaf’s top 1% of transcripts consumed 59.8% of all FL reads versus 49.2% in flower, leaving the remaining 99% of leaf transcripts competing for only 40.2% of sequencing depth. Per-condition analysis of the four Iso-Seq libraries confirmed that this skew is intrinsic to the leaf organ rather than an artefact of combining spring and summer reads before clustering: top-1% FL-read share was ≈ 55% in both leaf libraries (spring leaf; summer leaf) and ≈ 44% in both flower libraries (spring and summer), indicating that combining the two environmental conditions amplifies skew by ∼5 percentage points uniformly across both tissues but does not create any leaf-specific signature. The downstream consequence is visible in per-gene transcript distribution (Figure 5). The two organs differ sharply in per-gene isoform support. In flower, most genes carried multiple isoforms, giving Cogent the multi-isoform evidence its graph algorithm depends on. In leaf, the opposite held; per-gene isoform support was far lower (Figure 5). With little multi-isoform evidence per gene, most leaf genes fall below the level of support Cogent needs to reconstruct a locus correctly, so the algorithm instead merges sequence-similar but distinct genes. Below a critical threshold of multi-isoform input, Cogent’s graph algorithm misinterprets sequence similarity between distinct genes as alternative isoforms of the same locus, yielding the diagnostic “median rises after Cogent” signature observed in our leaf data (Figure 5D). Of two candidate input-data diagnostics, only one discriminated success from failure here. Median FL reads per transcript sets a necessary depth floor (≥ 4 typical) but was identical in the two tissues (4 in both) and so did not distinguish them; the top 1% FL-read concentration was the diagnostic that separated flower from leaf (49.2% versus 59.8%; suitable below ∼55%). These diagnostics can be computed from Iso-Seq HQ output before committing to graph reconstruction and would have predicted the leaf failure observed here. These thresholds are derived from a single species across two organs and should therefore be regarded as preliminary, indicative cutoffs rather than validated rules of thumb; cross-species and cross-tissue/organ validation will be required before they can be recommended as general predictors.

This pattern contrasts with the genome-free Iso-Seq workflow described by Li et al. (2017), who applied CD-HIT at 99% identity prior to Cogent reconstruction in *Astragalus membranaceus* and recovered transcriptome assemblies for both root and leaf tissues without the leaf-specific failure observed here. Three differences between the two studies plausibly explain this divergence. First, input scale forced our identity threshold. Li et al. processed Iso-Seq inputs of ≈ 115,725 (root) and 102,334 (leaf) combined HQ+LQ transcripts, at which scale CD-HIT 99% is a viable mild-redundancy filter. Our HQ-only input was an order of magnitude larger (1,150,020 flower; 803,912 leaf transcripts), and CD-HIT at 99% applied to input of this size would yield a post-filter set well above the ≤200,000-transcript ceiling recommended for direct Mash partitioning in the Cogent documentation; the 95% threshold was therefore selected to bring the input within Cogent’s tractable range. The unavoidable trade-off is that 95% collapses splice variants and near-paralogs lying between 95% and 99% identity, and in low-coverage tissues this collapse pushes per-gene multi-isoform support below the level required for reliable graph reconstruction. Second, *M. arvensis* is a paleopolyploid (Lysak et al., 2005) in origin but has since returned to a functionally diploid state, increasing the abundance of near-paralogs that CD-HIT 95% can collapse but Cogent then misattributes as single-locus isoforms; *A. membranaceus* is diploid, reducing the impact of this effect. Third, expression differences may play a role, as roots lack the photosynthetic gene dominance seen in leaves, and their leaf tissue may likely have had lower stress-response expression than ours. An additional confound is that our Iso-Seq sample for each condition × organ combination was a pool of five individuals, so allelic variation among individuals further inflates the population of near-identical sequences that CD-HIT collapses and Cogent then treats as single-locus structure. Together, these differences suggest that Cogent reconstruction is most likely to succeed when the pre-filter preserves per-gene multi-isoform diversity, the genome is diploid or near-diploid, and per-organ expression is not strongly skewed by hyper-expressed transcripts.

### Pipeline selection across organs

Evaluated against the three selection criteria (BUSCO completeness, unique mapping rate, and redundancy), the data support a different optimal pipeline for each organ. For flower, V1.c (Cogent + CD-HIT 98%) achieved 95.3% BUSCO completeness with an elevated single-copy proportion (Figure 3A, Table 1), 22.4 ± 1.5% unique mapping with 77.7 ± 2.4% total mapping rate (Figure 3B, Table 1), and substantial redundancy reduction relative to the IsoSeq HQ input (89,949 vs 1,150,020 transcripts; Table 1). All three criteria, therefore, favoured V1.c as the flower reference. For leaf, the same three criteria led to a different choice: V1.a (CD-HIT 95% direct, no Cogent) preserved BUSCO completeness at 90.1%, substantially higher than V1.b (78.0%), while delivering 86.82% total mapping rate (Figure 4B, Table 2). Unique mapping in leaf V1.a was lower than in flower V1.c (6.3% vs 11.9%), but for RSEM-based downstream analysis the expectation-maximisation algorithm probabilistically resolves multi-mapped reads, so total mapping rate is the primary determinant of expression-quantification accuracy. However, in leaf, identifying non-redundant transcripts required an alternative strategy based on expression-driven read clustering. Therefore, selecting Cogent for flower but omitting it for leaf reflects organ-specific data characteristics rather than methodological inconsistency. This highlights the importance of evaluating pipelines on an organ-by-organ basis rather than assuming a single optimisation is universally applicable.

## Conclusion

This study presents a complete genome-free Iso-Seq pipeline for non-model plant transcriptomics and demonstrates that pipeline performance varies across organs. For flower, Cogent graph reconstruction followed by CD-HIT at 98% identity (V1.c) yielded a high-quality reference transcriptome (95.3% BUSCO complete, 77.7 ± 2.4% total RSEM mapping). For leaf, by contrast, Cogent reconstruction proved detrimental; the best reference was obtained by redundancy removal alone, using CD-HIT at 95% identity with expression-based clustering (strategy V1.a) (90.1% BUSCO completeness, 86.82% total RSEM mapping). This divergence is driven by organ-specific full-length read distributions, showing that input-data characteristics determine the success of graph-based reconstruction. Several limitations bound the conclusions of this work. First, *M. arvensis* is paleopolyploid, and CD-HIT identity-based clustering at 95–98% thresholds cannot reliably distinguish recent paralog pairs from allelic variants, likely overestimating BUSCO duplication rates. Second, gene-level biological validation by multiple sequence alignment was limited to six benchmark genes without complex gene families not directly evaluated. Third, long Cogent-reconstructed paths (up to 17,572 bp) carry an unassessed chimaera risk. Additionally, we caution against using EvidentialGene (tr2aacds.pl) on Iso-Seq data. In our V2 pipeline, it reduced BUSCO completeness (from 94.3% to 92.7%) by discarding biologically valid full-length isoforms, showing that short-read filtering heuristics are not well suited for long reads. Future work should validate these references against an upcoming high-quality *M. arvensis* genome, test these organ-specific diagnostics across other tissues and species, and benchmark against nanopore platforms. Despite these limitations, this pipeline provides a robust, genome-free framework for full-length reference transcriptome generation in non-model species and offers guidance for future Iso-Seq projects.

## ACKNOWLEDGEMENTS

We are grateful to Gema Batanero, Ángel Caravantes, and Raquel Sánchez for technical support, and to Ángel Martín Alganza for his assistance with the computing facilities of the Research Unit ‘Modeling Nature’. This research was supported by grants from the Spanish Ministry of Science and Innovation (PID2021-126456NB and PID2025-168039NB, co-funded by EU FEDER funds). This work is a contribution to the Research Unit ‘Modelling Nature’, funded by the Consejería de Economía, Conocimiento, Empresas y Universidad (IE2017-5537 and QUALIFICA 00011), and to Evoflor (UGR), Unidad Asociada I+D+I al CSIC.

## Supplementary Material

**Suppl. Methodology I: *Gene-level clustering of the leaf reference***

To group the 130,407 leaf V1.a transcripts into genes and their isoforms without a reference genome, two complementary reads-clustering strategies were evaluated: expression-aware clustering with Corset (v1.09; Davidson and Oshlack, 2014), and sequence-based clustering with isONclust3 (Petri and Sahlin, 2025). For Corset, short-read RNA-seq libraries from leaf organ (four spring and four summer biological replicates; Gómez et al., 2020) were quasi-mapped to the V1.a transcripts using Salmon (v.1.5.2); Patro et al., 2017) with equivalence-class export (--dumpEq). Salmon equivalence classes, the groups of transcripts to which multi-mapping fragments are jointly assigned were supplied to Corset, which clusters transcripts by the proportion of shared multi-mapping reads together with their relative expression across samples, using a contig-ratio test to separate paralogues with divergent expression profiles. Replicates were assigned to two groups (spring, summer) via the -g option, and a minimum count of ten reads per transcript was applied (-m 10).

For isONclust3, the V1.a transcript FASTA was converted to FASTQ with uniform high (Q40) base-quality values and clustered in PacBio mode (default parameters: k = 15, w = 51, minimiser-sharing threshold = 0.5). Because isONclust3 was developed for error-containing long reads, applying it to polished transcripts with uniform quality uses it as a minimiser-based sequence clusterer with no expression component; this provides a sequence-only counterpart to the expression-aware Corset clustering on the identical transcript set.

To assess how faithfully each method preserved distinct genes, a representative set comprising the single longest transcript per cluster was extracted from the V1.a reference for each clustering, and its completeness assessed with BUSCO (v5.8.3; brassicales_odb10, n = 4,596) in transcriptome mode. These representative sets were used solely for method comparison; the complete V1.a transcript set was retained as the leaf reference, with the selected clustering supplying its gene-level grouping for annotation and quantification. Differential-expression quantification was performed independently with RSEM (v1.3.1) and Bowtie 2, consistent with the rest of the study; gene-level estimates were obtained by supplying the Corset cluster assignments to rsem-prepare-reference via --transcript-to-gene-map. Functional annotation (Trinotate; TransDecoder ORF prediction) was performed on the V1.a transcripts and collapsed to gene level using the same cluster assignments.

**Suppl. Figure 1.**
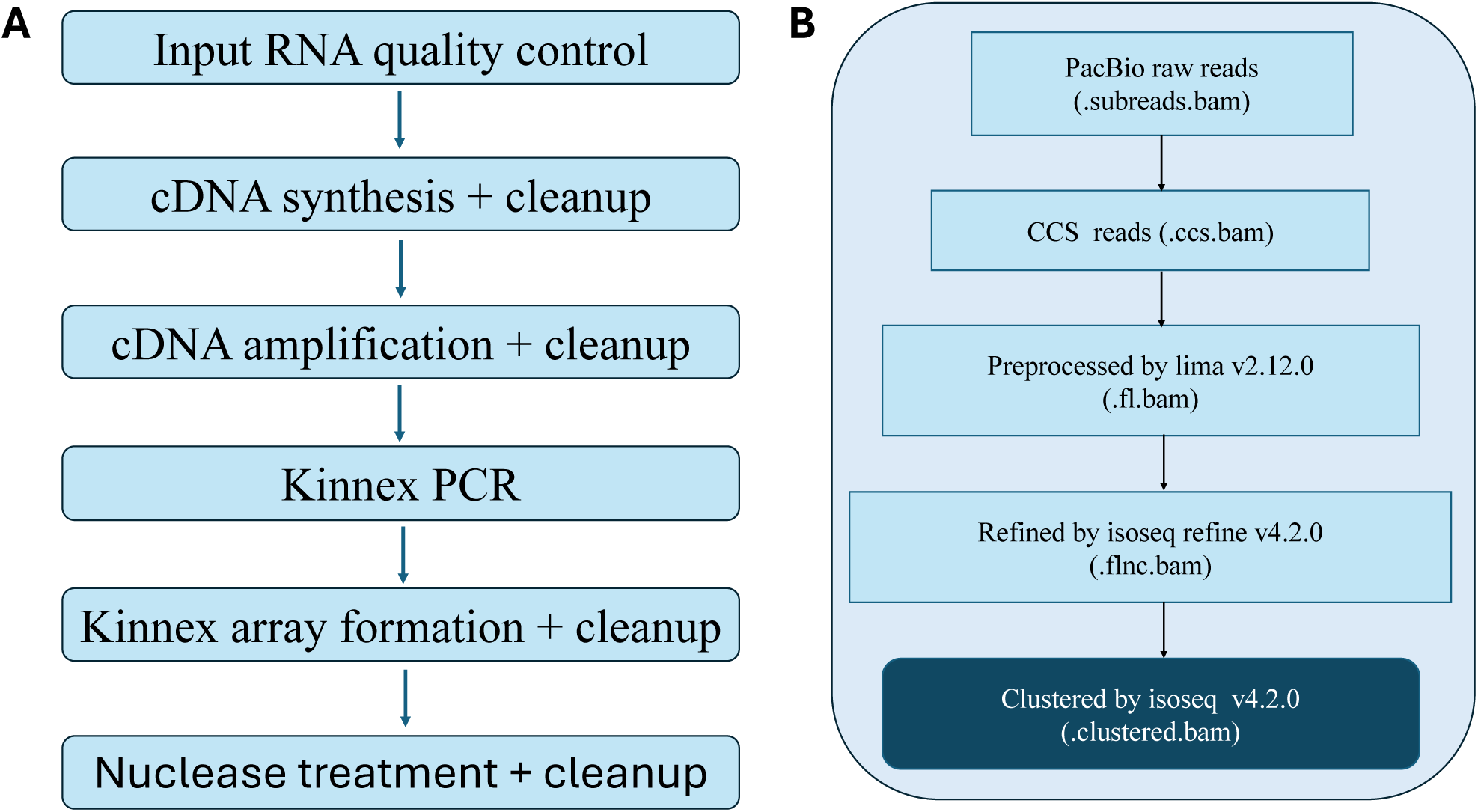
A) The Workflow of Kinnex Full-Length RNA Library Construction. B) Raw Iso-Seq reads were processed through the PacBio IsoSeq3 pipeline to obtain high-quality clustered transcripts.

**Suppl. Figure 2.**
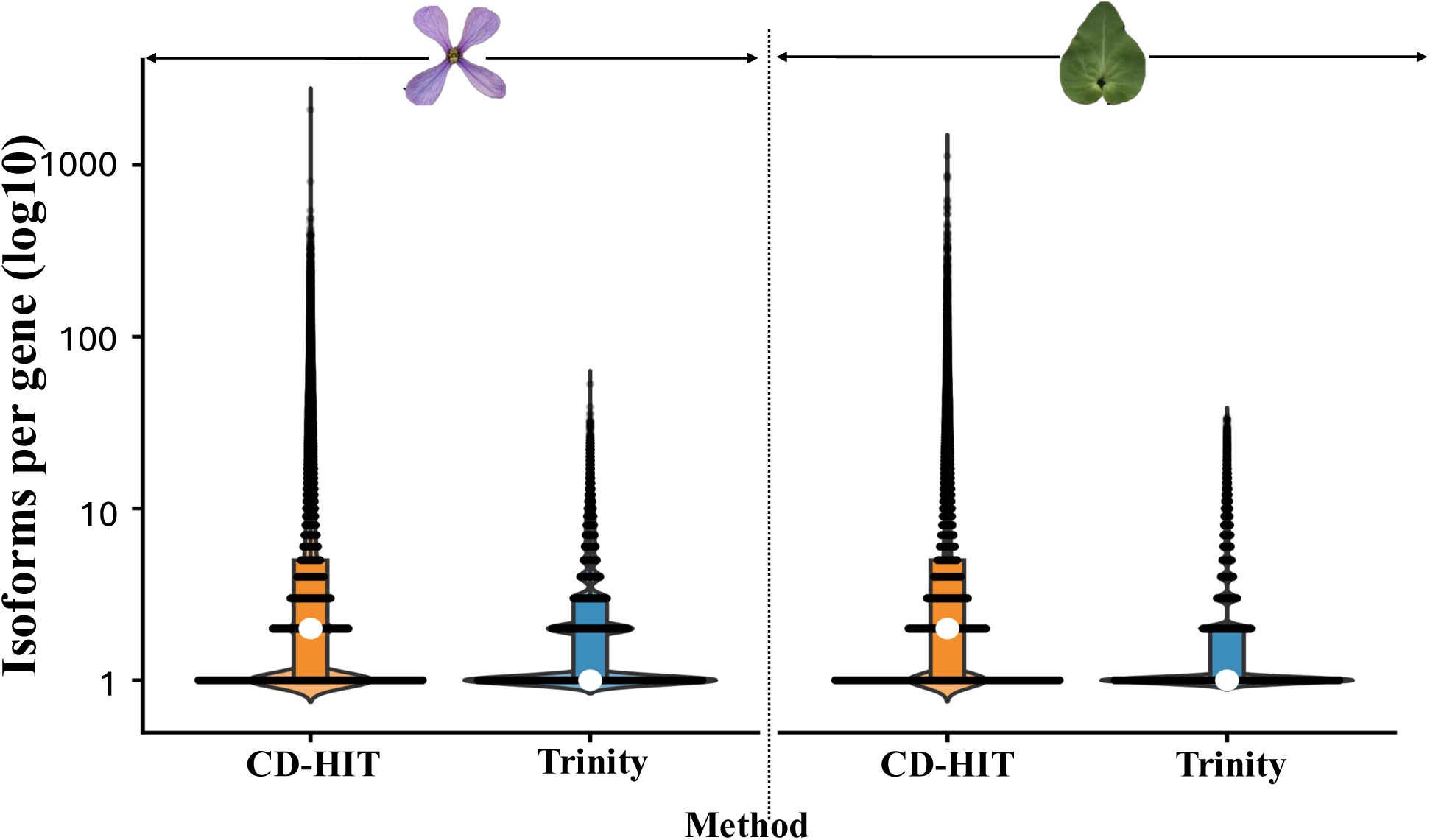
Number of isoforms per gene in CD-HIT 95% clustered Iso-Seq transcripts and the Trinity short-read assembly, illustrating residual redundancy at standard identity thresholds.

**Suppl. Figure 3.**
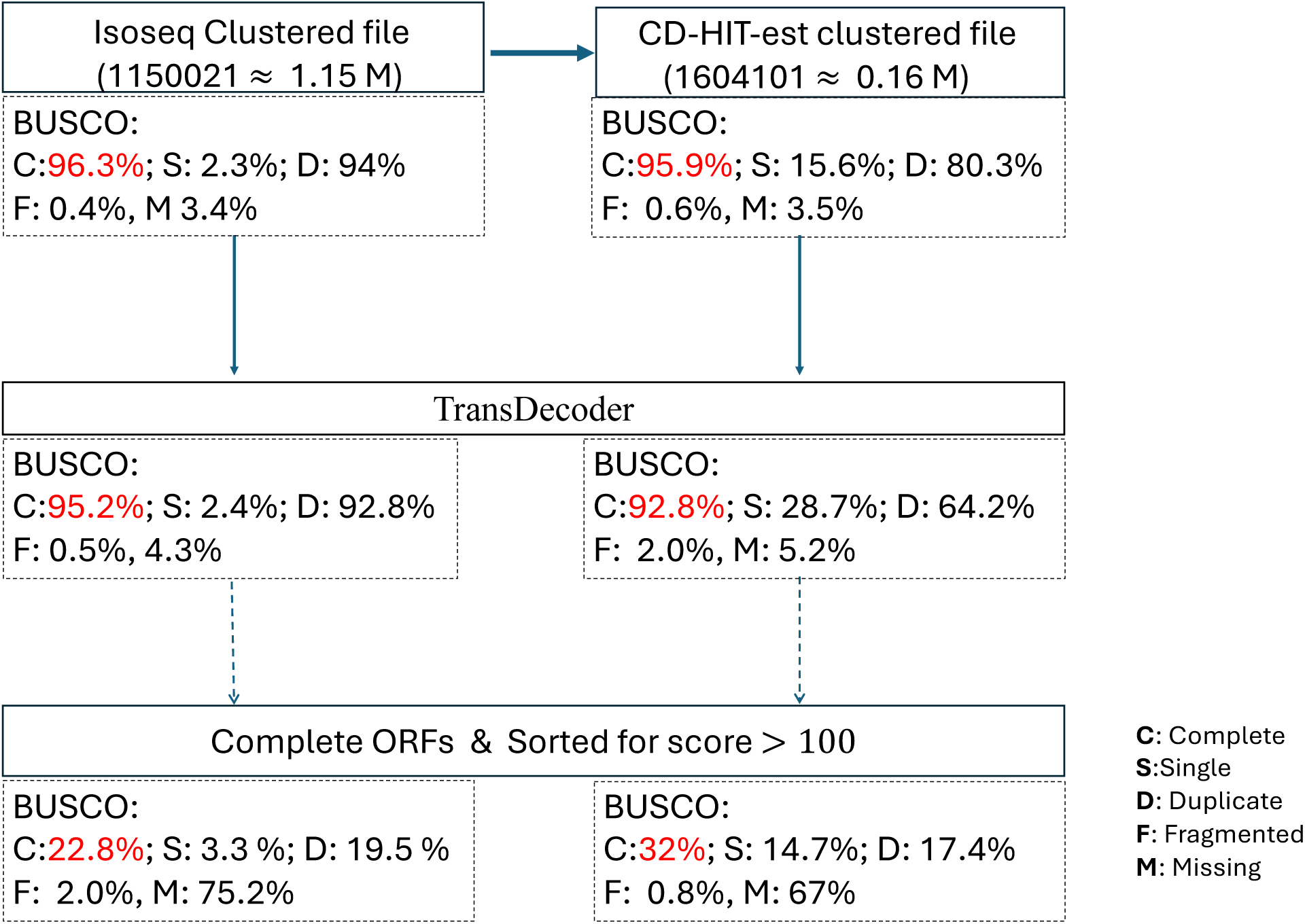
Additional BUSCO-based strategies for evaluating completeness of Iso-Seq de novo transcriptome assemblies in flower tissue.

**Suppl. Figure 4.**
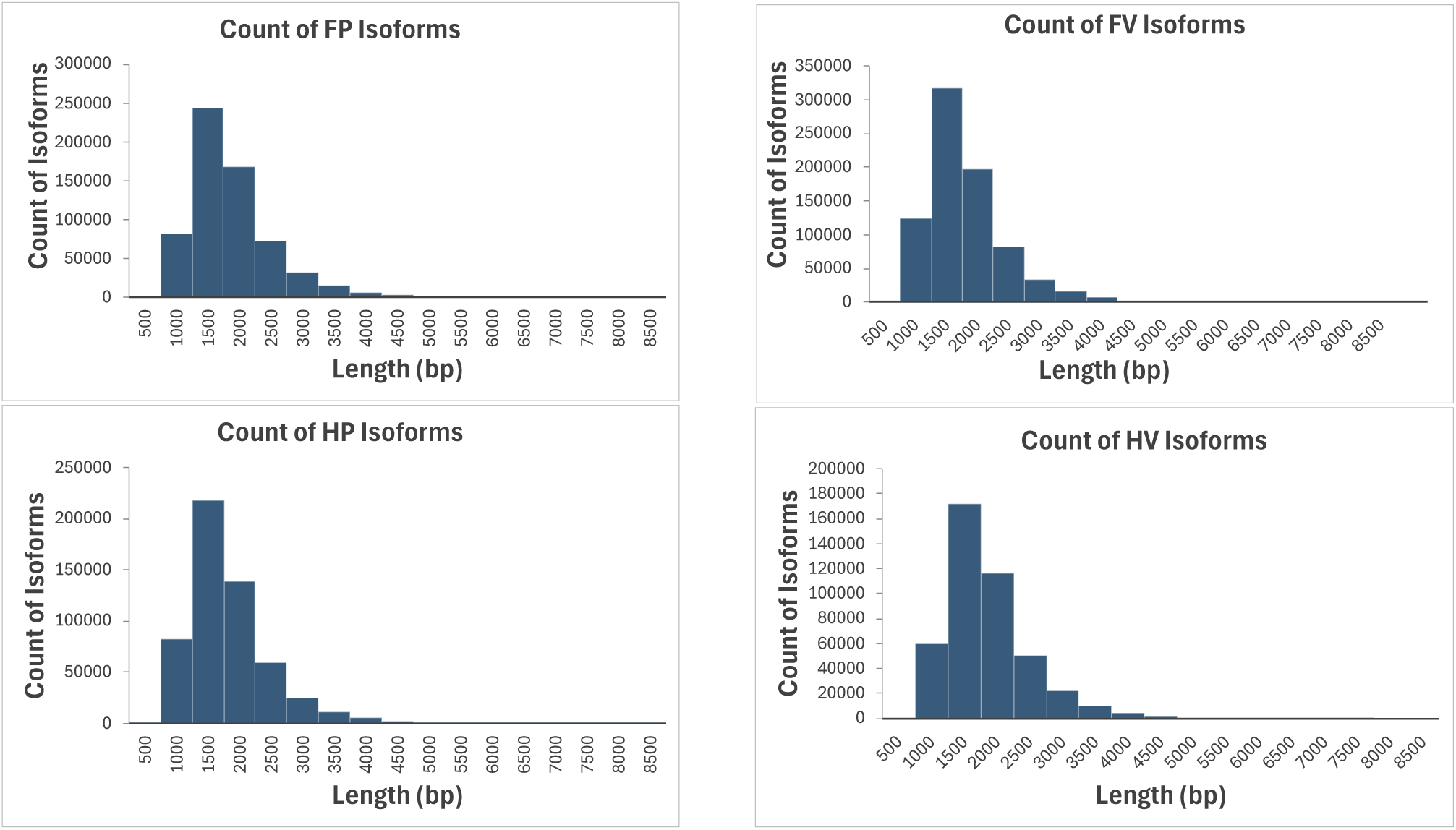
Count and length distribution of flower and leaf Iso-Seq isoforms across spring and summer conditions following Iso-Seq clustering. FP: spring flower; FV: summer flower; HP: spring leaf, HV: summer leaf.

**Suppl. Figure 5.**
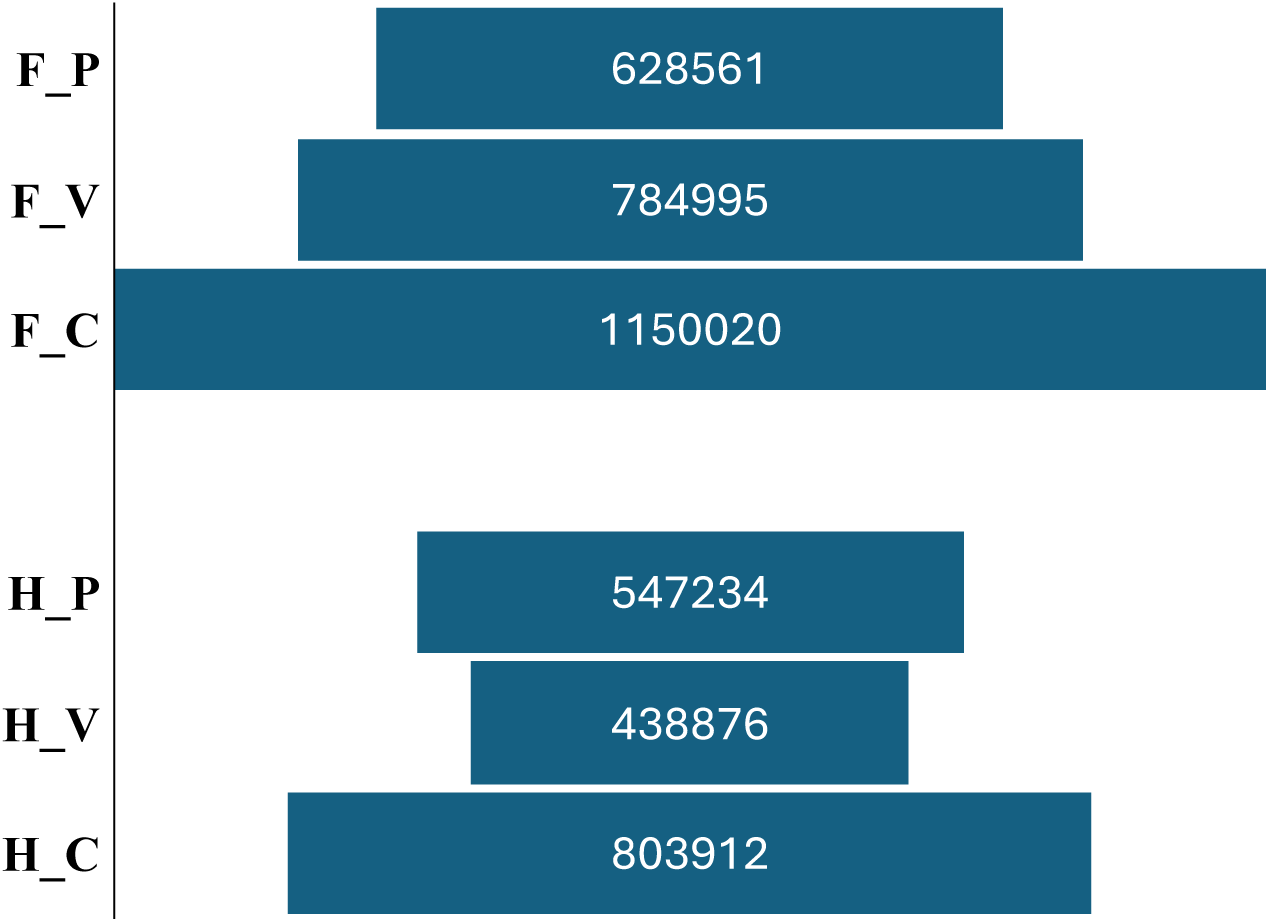
Iso-Seq cluster yield in combined and separated flower and leaf tissues across spring and summer conditions. F_P: spring flower; F_V: summer flower; H_P: spring leaf; H_V: summer leaf.

**Suppl. Figure 6.**
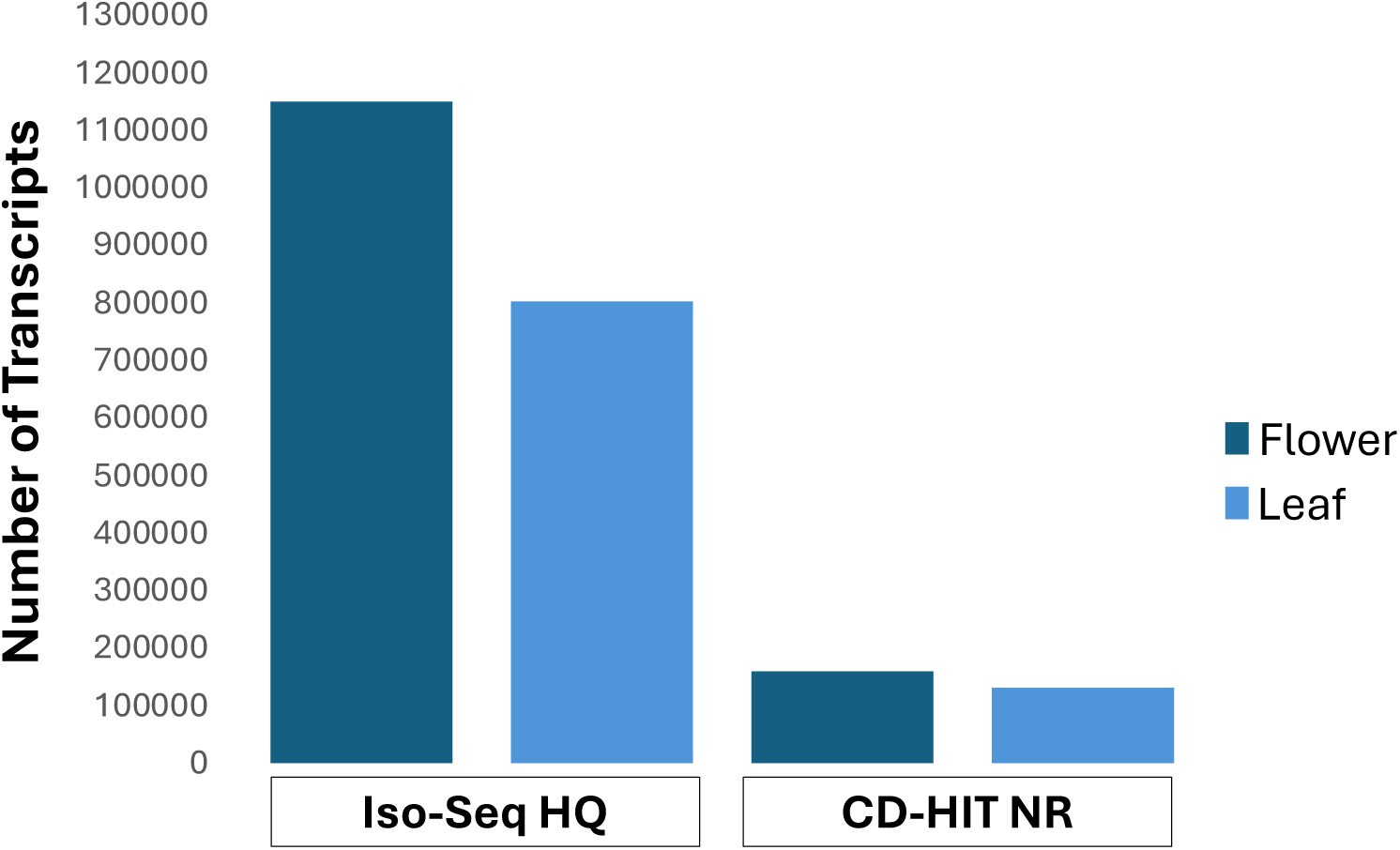
Transcript clustering and isoform diversity across the Iso-Seq pipeline. Total transcript counts at the high-quality Iso-Seq clustered (Iso-Seq HQ) and CD-HIT non-redundant (CD-HIT NR) stages for flower and leaf tissues.

**Suppl. Figure 7.**
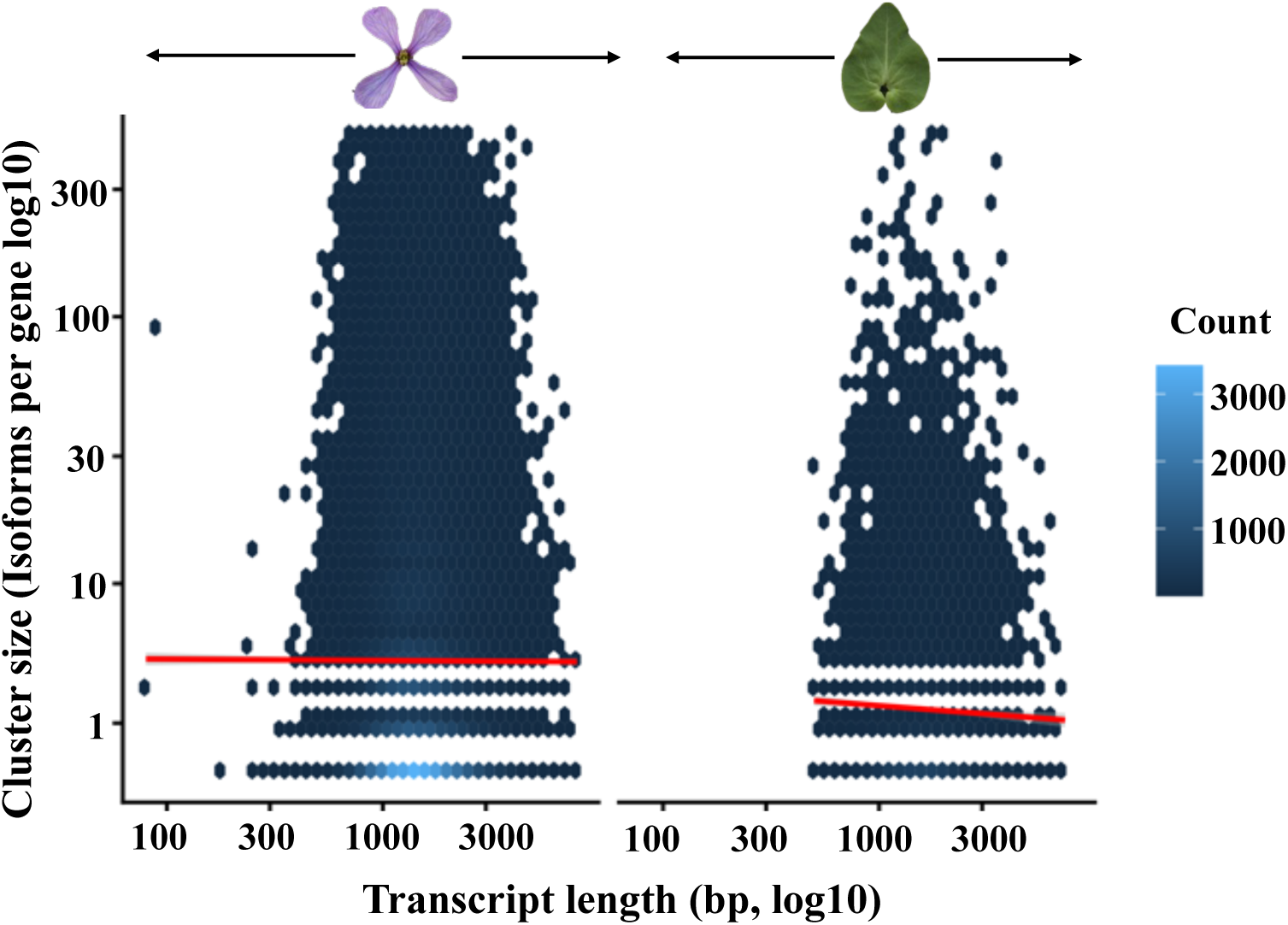
Mean number of isoforms per Cogent-reconstructed gene (or per CD-HIT 95% cluster) in flower and leaf tissues.

**Suppl. Figure 8.**
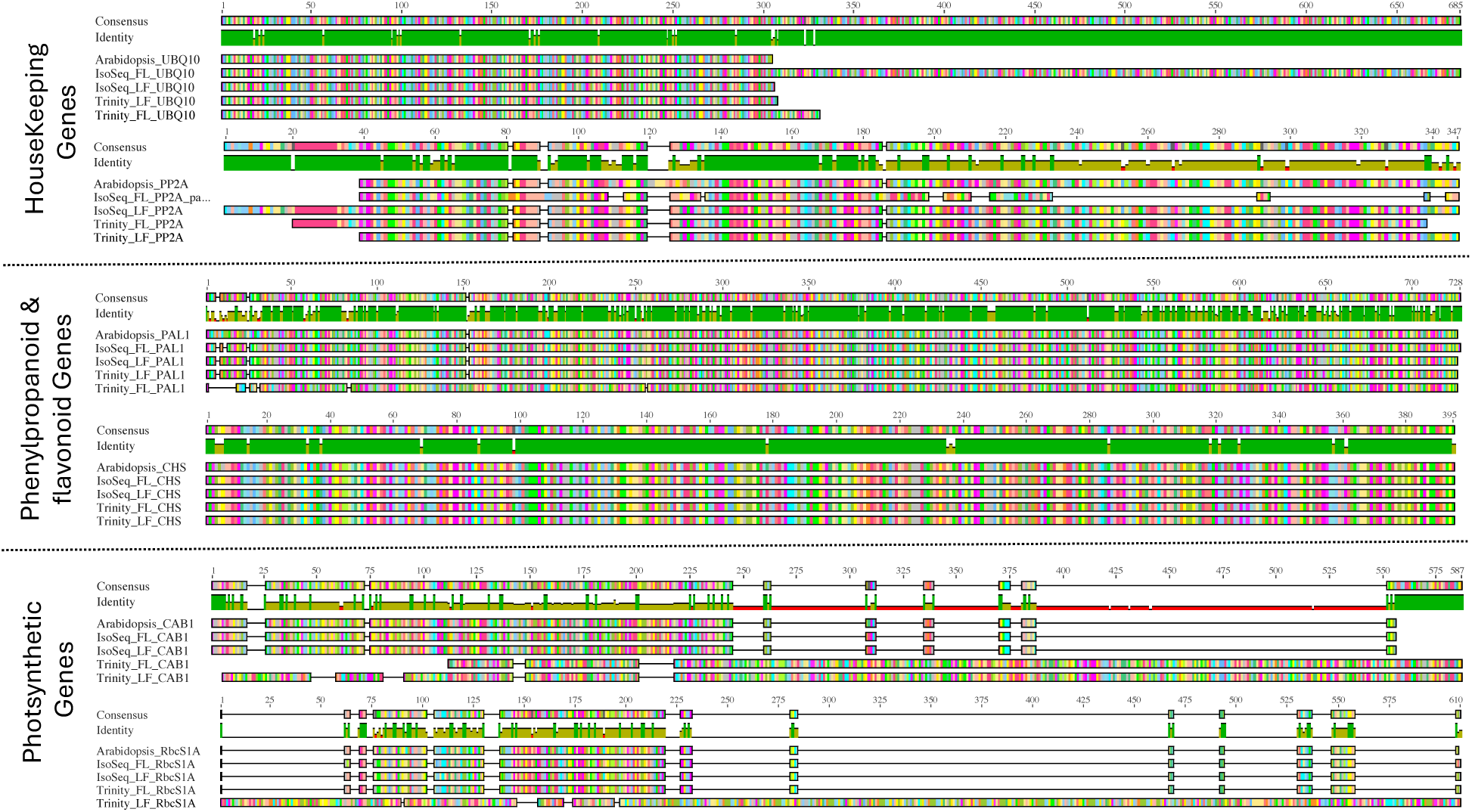
Multiple sequence alignment of predicted proteins from *Moricandia arvensis* Iso-Seq (flower and leaf) and Trinity (flower and leaf) references against *Arabidopsis thaliana* orthologues. Open reading frames were predicted using TransDecoder v5.5.0, aligned with MAFFT v7.562 (--auto --reorder), and visualized in Geneious Prime with percentage identity coloring relative to the Arabidopsis reference sequence (shown at top of each panel). IsoSeq_FL and IsoSeq_LF: Iso-Seq assembled flower and leaf transcripts, respectively. Trinity_FL and Trinity_LF: Trinity assembled flower and leaf transcripts, respectively.

**Suppl. Table 1.** CD-HIT-EST sequence clustering metrics at 98% and 95% identity thresholds for flower and leaf Iso-Seq HQ transcripts. Values shown for each identity threshold and tissue include total output clusters, collapse rate, isoform distribution statistics, and compatibility with Cogent’s recommended input ceiling (≤200,000 transcripts for direct Mash-based family finding). The 95% threshold was selected for downstream Cogent input based on two criteria: (1) output size within the Cogent-compatible range for both tissues, and (2) median cluster size of ≥2, indicating sufficient per-gene multi-isoform evidence for graph reconstruction.

| Metric | Flower |  | Leaf |  |
| --- | --- | --- | --- | --- |
|  | CD-HIT 98% | CD-HIT 95% | CD-HIT 98% | CD-HIT 95% |
| Total transcripts (input) | 1,148,231 | 1,148,231 | 803,912 | 803,912 |
| Total clusters (output) | 263,854 | 160,410 | 206,095 | 130,407 |
| Sequences collapsed (%) | 77.0% | 86.0% | 74.4% | 83.8% |
| Mean isoforms/gene | 4.36 | 7.16 | 3.90 | 6.16 |
| Median isoforms/gene | 1 | 2 | 2 | 2 |
| Min isoforms | 1 | 1 | 1 | 1 |
| Max isoforms | 1,805 | 2,092 | 425 | 1,126 |
| Singleton clusters (n, %) | 147,201 (55.8%) | 70,873 (44.2%) | 107,263 (52.0%) | 58,331 (44.7%) |
| Within Cogent limit ($\leq 200k$) | No | Yes | Yes | Yes |

**Suppl. Table 2.** Isoform distribution in CD-HIT 95% clustered transcripts and Cogent-reconstructed gene families for flower and leaf organs. Values are pre-cluster/per-family counts and are not directly comparable to the reconstructed-locus totals reported in the main text.

| Metric | Leaf CD-HIT 95% | Leaf Cogent | Flower CD-HIT 95% | Flower Cogent |
| --- | --- | --- | --- | --- |
| Total clusters/families | 130,407 | 22,450 | 160,410 | 34,075 |
| Total isoforms | 802,754 | 349,969 | 1,148,231 | 93,601 |
| Mean isoforms/gene | 6.16 | 5.34 | 7.16 | 2.75 |
| Median isoforms/gene | 2 | 4 (changed) | 2 | 2 (stable) |
| Min isoforms | 1 | 1 | 1 | 1 |
| Max isoforms | 1,126 | 90 | 2,092 | 111 |
| Singleton genes | 58,331 (44.7%) | 5,987 (9.1%) | 70,873 (44.2%) | 12,998 (38.1%) |

**Suppl. Table 3.** Length distribution metrics across processing stages for flower and leaf Iso-Seq data.

| Flower Transcriptome length statistics |  |  |  |  |  |  |  |
| --- | --- | --- | --- | --- | --- | --- | --- |
| Dataset | Sequences | Min (bp) | Max (bp) | Mean (bp) | Median (bp) | Total bases (bp) | N50 (bp) |
| IsoSeq HQ | 1,150,020 | 80 | 8,645 | 1,587 | 1,439 | 1,825,075,313 | 1,676 |
| + CD-HIT 95% | 160,410 | 500 | 8,645 | 1,666 | 1,531 | 267,201,677 | 1,777 |
| V1.b (Cogent raw) | 93,601 | 500 | 17,572 | 1,822 | 1,666 | 170,546,938 | 1,969 |
| V1.c (Cogent + CD-HIT 98%) | 89,949 | 500 | 17,572 | 1,819 | 1,662 | 163,646,272 | 1,967 |

| Leaf Transcriptome Flower Transcriptome length statistics |  |  |  |  |  |  |  |
| --- | --- | --- | --- | --- | --- | --- | --- |
| IsoSeq HQ | 803,912 | 71 | 8,224 | 1,593 | 1,444 | 1,280,261,210 | 1,682 |
| + CD-HIT 95% | 130,407 | 500 | 8,224 | 1,679 | 1,540 | 218,929,891 | 1,790 |
| Cogent reference (rejected) | 65,688 | 504 | 19,586 | 1,842 | 1,687 | 120,964,741 | 1,970 |

N50 is the length such that 50% of total assembled bases lie in transcripts of this length or longer. Length filter $\geq 500$ bp was applied prior to CD-HIT 95% pre-filtering for both tissues. Cogent v8.0.0 was run with default reconstruction parameters ( $k = 30$ ); the small additional length increase observed after CD-HIT 98% post-processing (V1.b → V1.c) reflects the removal of shorter near-identical isoforms.

**Suppl Table 4.**
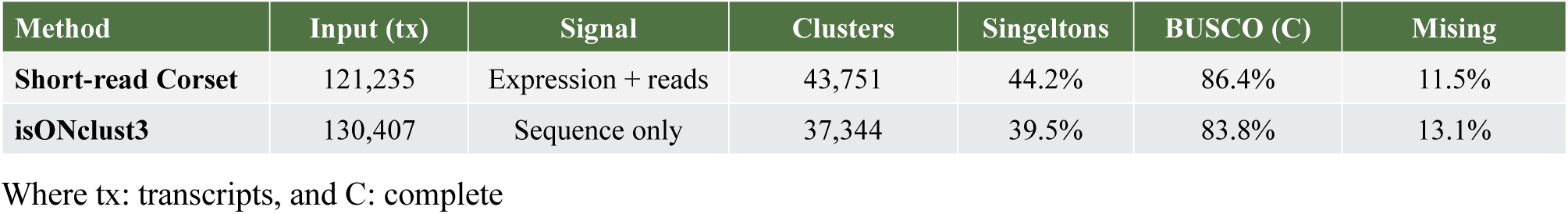
Comparison of leaf gene-clustering methods on the V1.a reference. Representative-per-cluster BUSCO (brassicales_odb10, n = 4,596) assessed on the longest transcript per cluster, extracted from V1.a. The complete V1.a transcript set (90.1%) was retained as the leaf reference; representative sets were used only for method comparison. Single-copy/duplicated split: short-read Corset 48.7/37.6; isONclust3 70.1/13.7; long-read Corset 37.3/27.1.

## REFERENCES

Ament, I.H., DeBruyne, N., Wang, F. and Lin, L., 2025. Long-read RNA sequencing: A transformative technology for exploring transcriptome complexity in human diseases. Molecular Therapy, 33(3), 883–894.

Bryant, D. M., Johnson, K., DiTommaso, T., Tickle, T., Couger, M. B., Payzin-Dogru, D., … & Whited, J. L. (2017). A tissue-mapped axolotl de novo transcriptome enables identification of limb regeneration factors. Cell reports, 18(3), 762–776.

Byrne, A., Cole, C., Volden, R. and Vollmers, C., 2019. Realizing the potential of full-length transcriptome sequencing. Philosophical Transactions of the Royal Society B, 374(1786), 20190097.

Camacho, C., Coulouris, G., Avagyan, V., Ma, N., Papadopoulos, J., Bealer, K., and Madden, T. L. (2009). BLAST+: architecture and applications. BMC Bioinformatics 10, 421. 10.1186/1471-2105-10-421

Clark, A., 2022. Developing omics and cell culture resources in non-model organism for their application in elucidating complex trait evolution in natural populations (Doctoral dissertation, Auburn University).

Csordás Tóth, É., Vissi, E., Kovács, I., Szöke, A., Ariño, J., Gergely, P., Dudits, D. and Dombrádi, V., 2000. Protein phosphatase 2A holoenzyme and its subunits from *Medicago sativa*. Plant Molecular Biology, 43, 527–536.

Cogent (v8.0.0). COding GENome reconstruction Tool. Available from Github: https://github.com/Magdoll/Cogent.

Cui, J., Shen, N., Lu, Z., Xu, G., Wang, Y. and Jin, B., 2020. Analysis and comprehensive comparison of PacBio and nanopore-based RNA sequencing of the *Arabidopsis* transcriptome. Plant Methods, 16(1), 85.

Davidson, N.M. and Oshlack, A., 2014. Corset: enabling differential gene expression analysis for de novo assembled transcriptomes. Genome Biology, 15(7), 410.

Fu, L., Niu, B., Zhu, Z., Wu, S. and Li, W., 2012. CD-HIT: accelerated for clustering the next-generation sequencing data. Bioinformatics, 28(23), 3150–3152.

Gilbert, Donald (2013) Gene-omes built from mRNA seq not genome DNA. 7th annual arthropod genomics symposium. Notre Dame. http://arthropods.eugenes.org/EvidentialGene/about/EvigeneRNA2013poster.pdf and http://globalhealth.nd.edu/7th-annual-arthropod-genomics-symposium/ doi:10.7490/f1000research.1112594.1

Gómez, J.M., Perfectti, F., Armas, C., Narbona, E., González-Megías, A., Navarro, L., DeSoto, L. and Torices, R., 2020. Within-individual phenotypic plasticity in flowers fosters pollination niche shift. Nature Communications, 11(1), 4019.

Grabherr, M. G., Haas, B. J., Yassour, M., Levin, J. Z., Thompson, D. A., Amit, I., … & Regev, A. (2011). Full-length transcriptome assembly from RNA-Seq data without a reference genome. Nature biotechnology, 29(7), 644–652.

Haas, B.J., Papanicolaou, A., Yassour, M., Grabherr, M., Blood, P.D., Bowden, J., Couger, M.B., Eccles, D., Li, B., Lieber, M., MacManes, M.D., Ott, M., Orvis, J., Pochet, N., Strozzi, F., Weeks, N., Westerman, R., William, T., Dewey, C.N., Henschel, R., LeDuc, R.D., Friedman, N. and Regev, A., 2013. De novo transcript sequence reconstruction from RNA-seq using the Trinity platform for reference generation and analysis. Nature Protocols, 8(8), 1494–1512.

Katoh, K, and Standley, D.M. (2013). MAFFT multiple sequence alignment software version 7: improvements in performance and usability. Molecular Biology and Evolution 30 (4), 772–780.

Kulaeva, O.A., Afonin, A.M., Zhernakov, A.I., Tikhonovich, I.A. and Zhukov, V.A., 2017. Transcriptomic studies in non-model plants: Case of *Pisum sativum* L. and *Medicago lupulina* L. In Marchi, F., Cirillo, P. and Mateo, E.C. (Eds.), Applications of RNA-Seq and omics strategies-from microorganisms to human health, 227–243. IntechOpen.

Langmead, B, and Salzberg, S. (2012). Fast gapped-read alignment with Bowtie 2. Nature Methods, 9: 357–359.

Li, B., Dewey, C.N., 2011. RSEM: accurate transcript quantification from RNA-Seq data with or without a reference genome. BMC Bioinformatics 12, 323. 10.1186/1471-2105-12-323

Li, J., Harata-Lee, Y., Denton, M.D., Feng, Q., Rathjen, J.R., Qu, Z. and Adelson, D.L., 2017. Long read reference genome-free reconstruction of a full-length transcriptome from *Astragalus membranaceus* reveals transcript variants involved in bioactive compound biosynthesis. Cell Discovery, 3, 17031.

Li, W. and Godzik, A., 2006. CD-HIT: a fast program for clustering and comparing large sets of protein or nucleotide sequences. Bioinformatics, 22(13), 1658–1659.

Lin, M.Y., Koppers, N., Denton, A., Schlüter, U. and Weber, A.P., 2021. Whole genome sequencing and assembly data of *Moricandia moricandioides* and *M. arvensis*. Data in Brief, 35, 106922.

Lockwood, B.L., Connor, K.M. and Gracey, A.Y., 2015. The environmentally tuned transcriptomes of *Mytilus* mussels. The Journal of Experimental Biology, 218(12), 1822–1833.

Lysak, M.A., Koch, M.A., Pecinka, A. and Schubert, I., 2005. Chromosome triplication found across the tribe Brassiceae. Genome Research, 15(4), 516–525.

Manni, M., Berkeley, M.R., Seppey, M. and Zdobnov, E.M., 2021. BUSCO: Assessing genomic data quality and beyond. Current Protocols, 1, e323.

Mante, J., Groover, K.E. and Pullen, R.M., 2025. Environmental community transcriptomics: strategies and struggles. Briefings in Functional Genomics, 24, elae033.

Manzoni, C., Kia, D.A., Vandrovcova, J., Hardy, J., Wood, N.W., Lewis, P.A. and Ferrari, R., 2018. Genome, transcriptome and proteome: the rise of omics data and their integration in biomedical sciences. Briefings in Bioinformatics, 19(2), 286–302.

Marks, R.A., Hotaling, S., Frandsen, P.B. and VanBuren, R., 2021. Representation and participation across 20 years of plant genome sequencing. Nature Plants, 7(12), 1571–1578.

Martin, J.A. and Wang, Z., 2011. Next-generation transcriptome assembly. Nature Reviews Genetics, 12(10), 671–682.

Máthé, C., Freytag, C., Kelemen, A., M-Hamvas, M. and Garda, T., 2023. “B” regulatory subunits of PP2A: their roles in plant development and stress reactions. International Journal of Molecular Sciences, 24(6), 5147.

Patro, R., Duggal, G., Love, M.I., Irizarry, R.A. and Kingsford, C., 2017. Salmon provides fast and bias-aware quantification of transcript expression. Nature Methods, 14(4), pp.417–419.

Petri, A.J. and Sahlin, K., 2025. De novo clustering of large, long-read transcriptome datasets with isONclust3. Bioinformatics, 41(5), btaf207.

Sharon, D., Tilgner, H., Grubert, F. and Snyder, M., 2013. A single-molecule long-read survey of the human transcriptome. Nature Biotechnology, 31(11), 1009–1014.

Shi, Z.X., Xiang, L., Zhao, H.M., Yang, L.Q., Chen, Z.C., Pu, Y.Q., Li, Y.W., Luo, B., Cai, Q.Y., Liu, B.L. and Feng, N.X., 2024. High-throughput single-molecule long-read RNA sequencing analysis of tissue-specific genes and isoforms in lettuce (*Lactuca sativa* L.). Communications Biology, 7(1), 920.

Soltis, P.S. and Soltis, D.E., 2021. Plant genomes: markers of evolutionary history and drivers of evolutionary change. *Plants, People*, Planet, 3(1), 74–82.

Steijger, T., Abril, J.F., Engström, P.G., et al. (2013). Assessment of transcript reconstruction methods for RNA-seq. Nature Methods, 10(12), 1177–1184. DOI: 10.1038/nmeth.2714

Wang, B., Tseng, E., Regulski, M., Clark, T.A., Hon, T., Jiao, Y., Lu, Z., Olson, A., Stein, J.C. and Ware, D., 2016. Unveiling the complexity of the maize transcriptome by single-molecule long-read sequencing. Nature Communications, 7(1), 11708.

Wang, Z., Gerstein, M. and Snyder, M., 2009. RNA-Seq: A revolutionary tool for transcriptomics. Nature Reviews Genetics, 10(1), 57–63.

Weirather, J.L., de Cesare, M., Wang, Y., Piazza, P., Sebastiano, V., Wang, X.J., Buck, D. and Au, K.F., 2017. Comprehensive comparison of Pacific Biosciences and Oxford Nanopore Technologies and their applications to transcriptome analysis. F1000Research, 6, 100.

